# Fluorescence Lifetime Imaging for Quantification of Targeted Drug Delivery in Varying Tumor Microenvironments

**DOI:** 10.1101/2024.01.12.575453

**Authors:** Amit Verma, Vikas Pandey, Catherine Sherry, Christopher James, Kailie Matteson, Jason T. Smith, Alena Rudkouskaya, Xavier Intes, Margarida Barroso

## Abstract

**Rationale:** Trastuzumab (TZM) is a monoclonal antibody that targets the human epidermal growth factor receptor (HER2) and is clinically used for the treatment of HER2-positive breast tumors. However, the tumor microenvironment can limit the access of TZM to the HER2 targets across the whole tumor and thereby compromise TZM’s therapeutic efficacy. An imaging methodology that can non-invasively quantify the binding of TZM-HER2, which is required for therapeutic action, and distribution within tumors with varying tumor microenvironments is much needed.

**Methods:** We performed near-infrared (NIR) fluorescence lifetime (FLI) Forster Resonance Energy Transfer (FRET) to measure TZM-HER2 binding, using *in vitro* microscopy and *in vivo* widefield macroscopy, in HER2 overexpressing breast and ovarian cancer cells and tumor xenografts, respectively. Immunohistochemistry was used to validate *in vivo* imaging results.

**Results:** NIR FLI FRET *in vitro* microscopy data show variations in intracellular distribution of bound TZM in HER2-positive breast AU565 and AU565 tumor-passaged XTM cell lines in comparison to SKOV-3 ovarian cancer cells. Macroscopy FLI (MFLI) FRET *in vivo* imaging data show that SKOV-3 tumors display reduced TZM binding compared to AU565 and XTM tumors, as validated by *ex vivo* immunohistochemistry. Moreover, AU565/XTM and SKOV-3 tumor xenografts display different amounts and distributions of TME components, such as collagen and vascularity. Therefore, these results suggest that SKOV-3 tumors are refractory to TZM delivery due to their disrupted vasculature and increased collagen content.

**Conclusion:** Our study demonstrates that FLI is a powerful analytical tool to monitor the delivery of antibody drug tumor both in cell cultures and in vivo live systems. Especially, MFLI FRET is a unique imaging modality that can directly quantify target engagement with potential to elucidate the role of the TME in drug delivery efficacy in intact live tumor xenografts.

## INTRODUCTION

Tumor microenvironment (TME) contributes to the impaired delivery of drugs and therapeutic resistance in breast cancer patients, leading to tumor relapse and metastasis [1–3]. A complex interplay occurs between cellular and acellular components of TME, such as tumor cells, immune cells, cytokines, fibroblasts, extracellular matrix, and blood vessels, to create dense, stiff, and refractory tumors. Tumor cells can modify TME leading to decreased drug delivery, penetration, exposure and drug-receptor engagement of antibody-based targeted therapeutic agents [2]. Trastuzumab (TZM) is a well-known humanized monoclonal antibody that targets human epidermal growth factor receptor (HER2) for the treatment of HER2-positive tumors. However, only 40 - 60 % of the patients respond to TZM therapy and the disease recurs in 15 - 20 % of breast cancer patients [1]. TME compromises the therapeutic efficacy of TZM by creating extracellular matrix and abnormal blood vessel barriers, which can restrict the distribution of TZM, and limit the access of TZM molecules to their targets [4]. Currently, there is a critical need to develop methods that can non-invasively quantify the amount of drug-target binding in live tissues and reveal the intra-tumoral distribution of antibody drug-target complexes in tumors with varying TME. Monitoring of TZM-HER2 drug-target engagement and efficacy in tumors with heterogenous TME will be instrumental to improving therapy response and efficacy, by the optimization of targeted delivery in breast cancer.

Immunohistochemical methods have been widely used to measure receptor expression as well as antibody-receptor binding in tumor tissue sections, but have limited dynamic range and multiplexing capacity, besides being an invasive procedure. Immuno-Positron Emission Tomography (PET) has been utilized for the quantification of drug-target engagement in various types of cancer such as breast, pancreatic, and ovarian cancer etc. [5–8]. However, immuno-PET uses maximum intensity projection to assess target expression, does not provide a direct and specific measurement of target-bound and unbound drug molecules, and cannot be easily multiplexed with other molecular and functional probes. Therefore, imaging preclinical approaches that allow for multiplexing distinct probes to monitor the molecular, metabolic, and functional characteristics of breast tumors before and after therapy are still lacking.

Optical imaging-based measurement of fluorescence decay to calculate the fluorescence lifetime (FLI) of near-infrared (NIR) fluorophores has been performed in breast cancer cells or tumor xenografts using preclinical wide-field macroscopy FLI (MFLI) imagers [9–12]. Importantly, NIR MFLI Forster Resonance Energy Transfer (FRET) measurements can quantify the fraction of drug binding to the target in live and intact preclinical animal models [13–15]. Thus, optical imaging has emerged as an invaluable tool to quantify the binding of antibody drugs to their target in a non-invasive and longitudinal manner [16,17]. We have demonstrated the utility of NIR MFLI FRET to measure receptor-ligand binding via dark quencher (QC-1)-mediated FRET in the transferrin (Tf) - transferrin receptor system [18]. QC-1 was used as a dark quencher FRET acceptor to prevent spectral bleed-through, increase the number of photons collected and provide higher sensitivity and specificity for the NIR MFLI FRET imaging performance. Recently, we successfully demonstrated MFLI FRET-based optical imaging to quantify the binding of TZM to its target receptor protein HER2, which is highly expressed in HER2-positive breast cancer cell lines [19,20]. The MFLI data indicates that the FRET signals, upon binding of NIR-labeled TZM FRET-pair probes to HER2 in intact, living HER2-positive tumor xenografts, can not only effectively delineate the tumor margins with high signal-to-noise ratio but also quantify the fraction of bound and internalized TZM-HER2 complexes [19,20].

Herein, we have subjected different HER2-positive tumor xenografts to MFLI FRET imaging using NIR-labeled TZM FRET probes. Importantly, these distinct HER2-positive tumor xenografts derived from breast AU565, XTM (derived from AU565 xenografts), and ovarian SKOV-3 cancer cells display clear differences in TME phenotypes. These three tumor types show distinct vascular bed morphology, collagen fiber content, and cancer cell vs. stroma ratio. These key TME parameters can influence the delivery of antibody drugs across the whole tumor volume and subsequently tumor response. Our results indicate that SKOV-3 tumors display reduced TZM binding compared to AU565 tumors as demonstrated via *in vivo* MFLI FRET imaging or *ex vivo* immunohistochemistry (IHC) analysis, although these two HER2 positive cancer cells possess similar levels of HER2 expression. SKOV-3 tumors display disrupted vasculature and increased collagen fiber content in contrast to AU565 tumors. Moreover, we have compared TZM-HER2 binding levels using MFLI FRET and IHC imaging across tumors ranging from the periphery to the center and showed a distinct distribution of tumor-bound TZM across whole AU565 vs SKOV-3 tumors. Our study demonstrates that the MFLI FRET imaging approach when applied to tumors displaying different TME phenotypes offers a powerful analytical tool to non-invasively monitor and quantify drug-target engagement in preclinical tumor models.

## EXPERIMENTAL METHODS

### Cell culture

HER2-positive AU565 and SKOV-3 ovarian cancer cell lines were obtained from ATCC. XTM is a cell line derived from cells isolated from AU565 xenograft. Breast cancer AU565 and XTM cells were maintained in RPMI 1640 medium supplemented with 10% fetal bovine serum, 10 mM HEPES, and 50 units/ml/ 50 µg/ml penicillin/streptomycin. SKOV-3 cells were maintained in McCoy’s media, supplemented with 10% fetal bovine serum and 50 units/ml/50 µg/ml penicillin/streptomycin. All the cell lines were incubated at 37^0^ C and 5 % CO_2_ and were used within passage 10 to prevent any changes in the phenotypic characteristics. For all item details refer to **Supplementary Table S1.**

### TZM labeling

TZM (anti-HER2 humanized monoclonal antibody) was conjugated to NIR probes donor (D) fluorophore (Alexa Fluor 700; AF700) or acceptor (A) fluorophore (Alexa Fluor 750; AF750) through monoreactive N-hydroxysuccinimide ester to lysine residues in the presence of 100 mM sodium bicarbonate, pH 8.3, according to manufacturer’s instructions. The probes were purified by Amicon Ultra-4-centrifugal filter units (MWCO 30 kDA;). The probes were washed extensively with PBS. The protein concentration was determined and adjusted to a concentration of 1 mg/ml followed by assessing the degree of labeling using spectrophotometer DU 640. The average degree of labeling was approximately 2 AF700 or AF750 dye molecules per TZM molecule. TZM-QC-1 conjugation was performed by Li-Cor with an average dye to protein ratio of 3. All ligands were normalized to concentration 1 mg/mL in phosphate-buffered saline pH 7.6 and filter sterilized.

### Labeled probe uptake and immunofluorescence

50,000 cells/well were plated on 8 well µ-slides and incubated at 37°C and 5% CO_2_ for 24 h. Then, cells were incubated with TZM-AF700 for another 24 h. After washing with PBS, cells were fixed with 4% paraformaldehyde and processed for immunostaining using mouse monoclonal anti-HER2 primary antibody (dilution 1:500), followed by secondary goat anti-mouse labeled with Alexa Fluor 488 (dilution 1:500). Photomicrographs were captured on Zeiss LSM 880 confocal microscope with identical settings across all the experiments and samples analyzed.

### Near-infrared fluorescence lifetime FRET microscopy

AU565, XTM, and SKOV-3 cells were plated on MatTek 35 mm glass bottom plates and after 24 h of cell growth at 37°C and 5% CO_2_, the cells were incubated with TZM-FRET pair probes: TZM-AF700 (Donor; D) or TZM-AF750 (Acceptor; A) for 24 h, using Acceptor to Donor (A:D) ratio of 0:1 (D: 20 µg/ml), 1:1 (A: 20 µg/ml & D: 20 µg/ml), 2:1 (A: 40 µg/ml & D: 20 µg/ml), and 3:1 (A: 60 µg/ml & D: 20 µg/ml). Cells were washed with Hank’s balanced basal salt solution (HBSS), fixed with 4% paraformaldehyde, and stored in DHB solution containing phenol red-free DMEM, 5 mg/ml BSA, 4mM L-glutamine, 20 mM HEPES, pH 7.4. NIR FRET time-correlated single photon counting (TCSPC) fluorescence lifetime microscopy (FLIM) was performed on a Zeiss LSM 880 Airyscan NLO multiphoton confocal microscope equipped with HPM-100-40 high-speed hybrid FLIM detector (GaAs 300-730 nm; Becker & Hickl) and a Titanium/ Sapphire laser (Ti: Sa) (680-1040 nm; Chameleon Ultra II,) as described in [16,19]. A Semrock FF01-716/40 bandpass filter and a Semrock FF01-715/LP blocking long-pass filter were inserted in the beam splitter assembly to detect the emission from AF700 and block scattered light, respectively. The 80/20 beam splitter in the internal beam splitter wheel in the LSM 880 was used to direct the 690 nm excitation light to the sample and to pass the emission fluorescence to the FLIM detector. Two-component exponential fitting using SPC Image software (Becker & Hickl; Berlin, Germany) was used to analyze the data and the χ^2^ fitness test was used to determine the validity of the fit, providing χ^2^ values.

### Animal Experiments

All the animal experiments were conducted with the approval of the Institutional Animal Care and Use Committee at Albany Medical College and Rensselaer Polytechnic Institute. Animal facilities at both institutions are accredited by the American Association for Accreditation for Laboratory Animal Care International. For generating tumor xenografts 10 x 10^6^ AU565, 10 x 10^6^ XTM, or 5 x 10^6^ SKOV-3 cells were mixed 1:1 with Cultrex BME and injected into the right and left inguinal mammary fat pad of 4-week-old athymic nude mice (CrTac: NCr-Foxn1nu;). The tumors were grown for 4 weeks and monitored daily. TZM FRET pair probes: TZM-AF700 (D) and TZM-AF750 (A) or TZM-QC-1 (dark quencher; A) were injected through retro-orbital route with A:D ratio of 2:1 (A: 40 µg/ml & D: 20 µg/ml), or 3:1 (A: 60 µg/ml & D: 20 µg/ml) in a 100 – 120 µl volume. We performed MFLI imaging of TZM-FRET pair probes post-injection (p.i.) in mice bearing AU565, XTM, and SKOV-3 tumor xenografts. **Figure S1** shows the injection and imaging protocols for AU565, XTM, and SKOV-3 tumor xenografts-bearing mice: **Protocol 1a (P-1a)**, mice bearing AU565 and SKOV-3 tumor xenografts were injected intravenously with TZM-AF700/TZM-AF750 (A:D 2:1) and live intact animal MFLI data was captured at 24 h and 48 h p.i. (**Figure S1A**); **Protocol-1b (P-1b),** mice bearing AU565 and SKOV-3 tumor xenografts were injected intravenously with TZM-AF700/TZM-QC-1 (A:D 3:1) and live intact animal MFLI data was captured at 24 h and 48 h p.i. (**Figure S1B**); **Protocol 2 (P-2)**, mice bearing XTM and SKOV-3 tumor xenografts were injected intravenously with TZM-AF700/TZM-QC-1 (A:D 2:1) and live intact animal MFLI data was captured at 24 h and 48 h p.i. (**Figure S1C**); **Protocol-3 (P-3)**, mice bearing AU565 and XTM tumor xenografts were injected intravenously with TZM-AF700/TZM-QC-1 (A:D 3:1) and live intact animal MFLI data was captured at 24 h and 48 h p.i. (**Figure S1D**).

### Wide-field Macroscopic Fluorescence Lifetime Imaging (MFLI) Platform

The wide-field time-domain fluorescence lifetime imaging system, the detailed description of the set-up in [21], was used to perform MFLI with slight modifications as per the experimental requirements. A tunable Ti-Sapphire laser was used as the excitation source (spectral range 690-1040 nm) with 100-fs pulses at 80 MHz, coupled to a digital micromirror (DMD) device for wide-field illumination over an 8 cm x 6 cm area with 256 grayscale levels encoding at each pixel. The modulation of wide-field illumination was achieved by controlling DMD via Microsoft PowerPoint to ensure optimal signal detection over the whole animal body [22]. A time-gated intensified CCD (ICCD) camera with a gate width of 300 ps was used to collect the time-resolved data. The Instrument Response Function (IRF) and fluorescence signals were collected with a gate delay of 40 ps over a 7.0 ns time window in reflectance set-up. The total number of gates acquired was 200, and the maximum range detection was 4096 photons per pixel per gate. The microchannel plate (MCP) employed for signal amplification was set to 510 V for fluorescence lifetime imaging. The laser excitation for AF700 was set at 700 nm and emission filters were 720 ± 6.5 nm (FF01-720/13-25, Semrock, IL, Rochester, NY, USA) and 715 nm long pass filter (Semrock, FF01-715/LP-25). The IRF was collected using a full field pattern projected from DMD on diffuse white paper, and the time-resolved images were captured with ND filter. For small animal imaging, the imaging platform was equipped with an isoflurane anesthesia machine and a digitally controlled warming pad, as described in [11,18]. The animals were imaged at 12 h, and 24 h p.i. in the above-described MFLI platform with consistent imaging parameters and conditions.

### Bi-exponential fitting to extract FRET donor fraction using MFLI imager *in vivo*

The FRET donor fraction (FD%) which indicates the fraction of the population of TZM-AF700 (D) in the proximity of TZM-QC-1 or TZM-AF750 (A), permitting FRET to occur (FRET range, 2-10 nm) within the region of interest (ROI) was calculated by fitting the fluorescence decays in each pixel of ROI to the bi-exponential model:

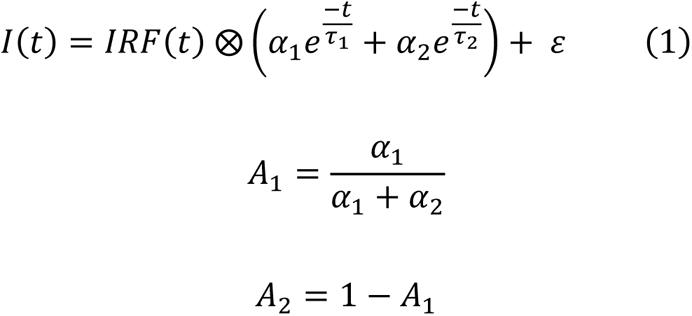

Here, I(t) represent observed fluorescence decay, IRF(t) is the instrument response function inherent to the system, α_1_ and α_2_ are the amplitude of the exponential terms; A_1_ and A_2_ are the FRET donor fraction (FD%) and non-FRET donor fractions respectively, τ_1_ and τ_2_ are the quenched and unquenched donor lifetimes, t is time, 𝜀 is an additive Poissonian noise, and ⊗ represents convolution operation. The temporal point spread function (TPSF) at each pixel was fitted using the open-source software named AlliGator [23] which uses the Levenberg-Marquardt non-linear least square fitting algorithm to estimate the optimized bi-exponential model parameters. The guess fitting parameter ranges for τ_1_ and τ_2_ were kept [0.2 – 0.6] ns and [0.9 – 1.5] ns respectively. The fluorescence decays from the MFLI system were analyzed with these fitting parameters and smoothing with Anscombe filtering.

### Immunohistochemistry (IHC)

All the tumors were extracted, fixed in 10% formalin and paraffin-embedded, followed by immunohistochemistry. The 5 µm tumor tissue sections were deparaffinized, rehydrated, and subjected to epitope retrieval using 1 mM EDTA pH 8.0 for 30 min. Vectastain ABC Elite kit was used for IHC staining with Vectro NovaRED as a peroxidase substrate (**Table S1**). Methyl Green was used for counterstaining the tumor tissue sections (**Table S1**). Hematoxylin & Eosin (H&E) stain was used for morphological characterization and Masson’s Trichrome staining was performed for staining collagen. The tumor tissue was stained for anti-HER2 (dilution 1:800), anti-CD31 (dilution 1:100), and rabbit monoclonal anti-TZM (dilution 1:100) primary antibodies (**Table S1**). Brightfield photomicrographs were captured using an Olympus BX40 microscope equipped with an Infinity 3 camera.

### Statistical Analysis

Unpaired Student’s t-tests and one way ANOVA was used for experimental data statistical analysis. Data are presented as mean with a 95% confidence interval or standard deviation as indicated. p-values less than 0.05 were considered statistically significant. Data were analyzed using MATLAB, Origin lab 8.0 or Microsoft Excel, and data visualization was generated using the PlotsOfData web app.

## RESULTS AND DISCUSSION

### Intracellular distribution of TZM-AF700 in HER2-positive human breast and ovarian cancer cell lines

The heterogeneity of TZM intracellular distribution has been shown previously in HER2-positive breast cancer and ovarian carcinoma cells [4,19,24,25]. Herein, we performed a comparative analysis of the intracellular distribution of TZM and HER2 in different human breast (AU565 and XTM) and ovarian (SKOV-3) HER2-positive cell lines, which vary in their morphological and physiological nature as well as tumor aggressiveness [4]. XTM are AU565-tumor passaged cells that maintain their HER2-positive nature AU565, XTM, and SKOV-3 cells were subjected to TZM-AF700 uptake for 24 h followed by fixation and HER2 immunostaining. Confocal microscopy images show that colocalization of labeled TZM-AF700 (red) with HER2 (green) occurs predominantly at the plasma membrane in AU565 and XTM cells compared to cytoplasmic/ perinuclear regions in SKOV-3 cells after 24 h post-TZM treatment (**Figure 1A**). However, no quantitative difference is detected in TZM-HER2 colocalization as shown by Pearson’s correlation coefficient measurements (**Figure 1B**). Immunoblotting data shows no significant difference in the HER2 protein expression levels among all three cell lines (**Figure 1C**). These results suggest increased intracellular accumulation of TZM-HER2 complexes in SKOV-3, in comparison to AU565 and XTM cells. Variations in the intracellular distribution of TZM may be an indicator of differential cellular responses to TZM treatment. However, colocalization of HER2 immunostaining with labeled internalized TZM-AF700 does not conclusively determine the fraction of HER2-bound TZM. Therefore, we have performed NIR FLIM FRET microscopy to quantify the fraction of TZM binding to HER2 in AU565 and SKOV-3 cells.

**Figure 1.**
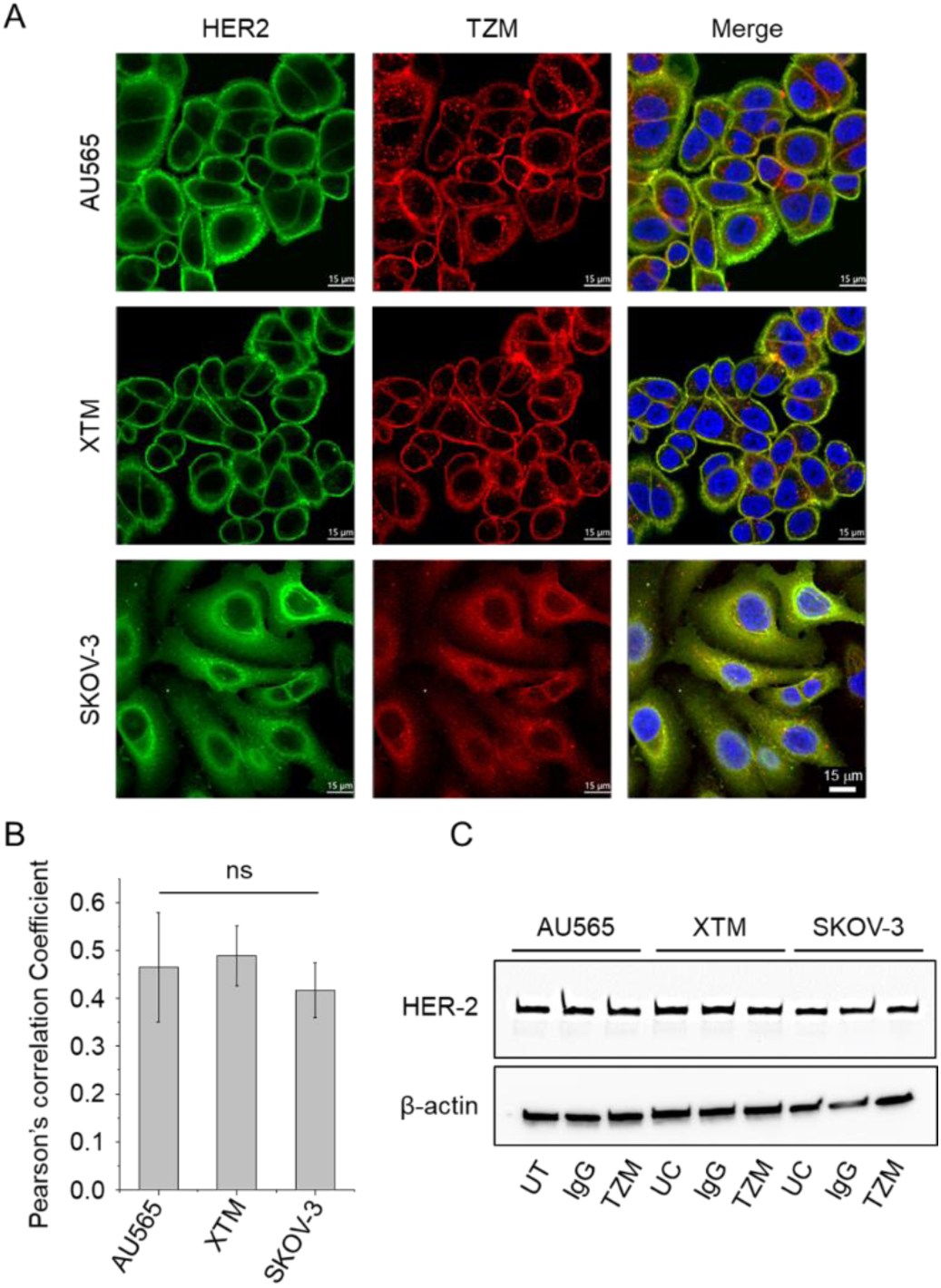
Intracellular distribution of HER2 and TZM complexes in HER2-positive breast (AU565 and XTM) and ovarian (SKOV3) cancer cell lines. **(A)** Photomicrographs show distribution of HER2 and TZM in AU565, XTM, and SKOV-3 cells. Immunofluorescence analysis of HER2 (green) and TZM (red) was performed in cells incubated with 20 µg/mL TZM–AF700 (red) for 24 h. Middle slices of z-stacks consisting of 6–8 optical slices are shown. Nuclei are visualized with DAPI. Scale bar = 15 µm. **(B)** Quantification of HER2 and TZM colocalization analysis using Pearson’s correlation coefficient across the z-stack. Imaris image analysis software was used for quantification of HER2-TZM colocalization. The bar graph shows the mean of five independent confocal z-stacks, n=5. Error bars indicate standard deviation. One-Way ANOVA for statistical analysis of colocalization between HER2 and TZM and p > 0.05 was considered not significant (ns). **(C)** Immunoblotting analysis of HER2 levels in whole cell lysates of AU565, XTM, and SKOV-3 incubated with IgG or TZM or without (UT) and probed with anti-HER2. Anti-β-actin was used as a loading control. All the protein samples were immunoblotted in triplicates.

### Quantification of TZM-HER2 drug-target engagement *in vitro*

Since differences in the endocytic and HER2 trafficking pathways can modulate TZM efficacy, it is important to characterize the intracellular distribution of TZM-HER2 complexes in several HER2-positive human cancer cell lines [26–28]. *In vitro* NIR FLIM FRET microscopy provides molecular information on the HER2-TZM binding and uptake into cancer cells [11,29,30]. In the present study, we investigated NIR FLIM FRET in AU565 cells and SKOV-3 cells, 24 h post incubation with TZM-AF700 alone or TZM-AF700 and TZM-AF750 FRET pair at different 0:1, 1:1, 2:1, and 3:1 A:D ratios. **Figure 2A** shows donor fluorescence intensity and lifetime (τ_m_) TCSPC images in the presence or absence of acceptor and at varying A:D ratios in AU565 and SKOV-3 cells. The donor fluorescence intensity data shows a differential distribution of labeled TZM in these cell lines, with AU565 cells showing TZM primarily at the plasma membrane and SKOV-3 cells displaying TZM mostly in cytoplasmic vesicles (**Figure 2A**). These results corroborate the immunofluorescence data in **Figure 1A**. The donor lifetime (τ_m_) images indicate a substantial reduction in donor lifetime with increasing A:D ratio, suggesting the occurrence of FRET at the plasma membrane and in intracellular vesicles in both AU565 and SKOV-3 cells (**Figure 2B**). The pixel-wise frequency distribution of donor lifetime data using bi-exponential fitting, the Levenberg-Marquardt algorithm as described in Eqn (1), with floating short and long lifetime components suggests that, as expected, donor lifetime decreases with increasing A:D ratio in AU565 cells and SKOV-3 cells (**Figure 2B-C**). Interestingly, we have detected variation in the average donor long lifetime component between AU565 and SKOV-3, probably due to their distinct intracellular localization of HER2-TZM complexes. Previously, we have shown that TZM-AF700 does not show fluorescence lifetime changes upon changes in pH [19,20]. Nevertheless, these results suggest TZM-AF700 can undergo lifetime changes as it undergoes endocytic internalization in these cells, probably due to alterations in their surrounding environment.

**Figure 2.**
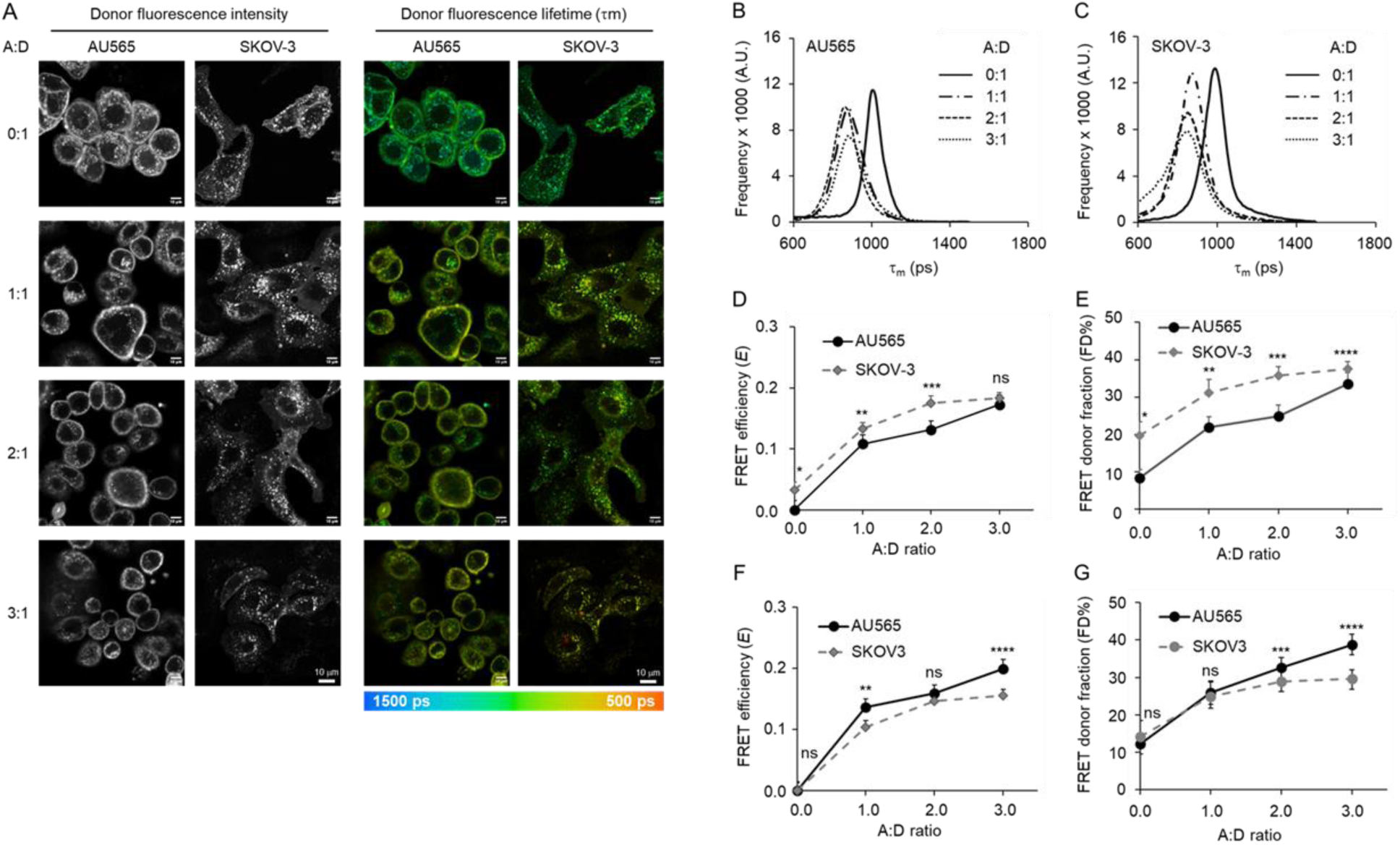
NIR FLI FRET microscopy in HER2 positive AU565 and SKOV-3 cells. **(A)** Fluorescence intensity and mean lifetime map (τ_m_) images of time-correlated single photon counting (TCSPC) in AU565 and SKOV-3 cells, treated with TZM-AF700 (Donor) or TZM–AF700 plus TZM–AF750 (Acceptor) with varying ratios Acceptor/Donor (A:D) = 0:1; 1:1, 2:1; 3:1; pseudo-color range = 500–1500 ps. Scale bar = 10 µm. Fluorescent lifetime distribution in **(B)** AU565 and **(C)** SKOV-3 cells treated with TZM–AF700 (A:D = 0:1) or TZM–AF700 and TZM–AF750 (A:D = 1:1, 2:1; 3:1). Comparison of TZM–FRET efficiency (*E*) **(D)** and FRET donor fraction (FD%) **(E)** calculated using fluorescence long lifetimes of TZM-AF700 extracted after averaging across both AU565 and SKOV-3 cells with varying A:D ratios (0:1, 1:1, 2:1, 3:1). Comparison of TZM–FRET efficiency (*E*) **(F)** and FRET donor fraction (FD%) **(G)** calculated using fluorescence long lifetimes extracted from each cell line (AU565 and SKOV-3 cells) of TZM-AF700 with varying A:D ratios (0:1, 1:1, 2:1, 3:1). Analysis was performed using 10 distinct pixel coordinates (n = 10) from five independent ROIs; error bars represent confidence interval at 95%. Data presented as mean ±confidence interval at 95%, n = 10. Asterisks indicate p<0.05 (significant); **Supplementary Table S2**.

Herein, we analyzed the NIR FLIM microscopy data in two distinct ways. Firstly, the long lifetime component was averaged across both AU565 and SKOV-3 single donor samples (A:D ratio = 0:1) and fixed to extract the short lifetime using bi-exponential fitting and calculate energy transfer efficiency (*E)* and the amplitude of the short lifetime, i.e., FD% (**Figure 2D-E**). Alternatively, the long lifetime component was extracted from each cell line single donor (A:D ratio = 0:1) samples and fixed to extract the short lifetime component using bi-exponential fitting and calculate *E* and FD% values (**Figure 2F-G**). As shown in **Figure 2D-G**, a significant increase in FRET efficiency, *E,* (**D & F**), and FD% (**E & G**) levels was observed in AU565 cells and SKOV-3 cells with increasing A:D ratios, indicating that binding and internalization of TZM-HER2 complexes occurs in both cell lines. Averaging the long lifetime leads to slightly increased *E* and FD% levels in SKOV-3 vs. AU565 cells, although the average FD% at A:D ratio = 0 (negative control) is significantly higher in SKOV-3 cells. Using cell-specific long lifetimes in FLIM TCSPC results in similar *E* and FD% levels for AU565 and SKOV-3 cells, suggesting that TZM-HER2 drug-target engagement occurs at comparable levels in both cell lines.

Overall, our results indicate that different HER2-positive cancer cell lines can present distinct intracellular distributions. In SKOV-3 cells, the ratio between HER2-TZM at plasma membrane vs. intracellular vesicles is altered, with more HER2-TZM complexes located intracellularly [19,20]. Nevertheless, SKOV-3 and AU565 show similar *E* and FD% levels, indicating that SKOV-3 cells can still bind and internalize TZM in an HER2-dependent manner. In summary, NIR FLIM FRET methodology can be robustly used to evaluate TZM-HER2 drug-target engagement, opening new avenues for advancing our understanding of TZM uptake and intracellular trafficking in HER2 positive cancer cell lines with distinct HER2 intracellular distributions.

### Morphological and molecular characterization of untreated and TZM-treated HER2- positive human breast and ovarian tumor xenografts

Drug response is driven directly by oncogenic receptor expression and mutational burden levels in tumor cells and modulated via various cellular/non-cellular components of TME, which can regulate drug distribution, penetration, and binding across solid tumors [31]. We performed morphological and molecular characterization of untreated and TZM-treated HER2 positive human breast (AU565 and XTM) and ovarian and (SKOV-3) tumor xenografts in mice using H&E staining and immunohistochemistry. The goal is to establish tumor xenograft mice models that vary in their TME and test the impact of these TME differences in drug-target engagement using MFLI FRET *in vivo* imaging.

After four weeks of tumor growth, nude mice were retro-orbitally injected with 60 µg of TZM (equivalent dose to NIR-labeled TZM injection in MFLI FRET imaging). Tumors were extracted 48 h post-treatment and subjected to sectioning for histopathological analysis. Post-TZM treatment, the H&E staining shows a decrease in cellularity in AU565 and XTM tumors compared to that in SKOV-3, suggesting that TZM induces elevated cytotoxicity in HER2-positive breast tumor xenografts in comparison to ovarian tumor xenografts (**Figure 3A, H&E**). Imaging doses of TZM (60 µg total) are significantly less than those used for therapeutic purposes (2 to 10 mg/ml) [32], warranting the reduced cytotoxicity levels. Masson’s Trichrome staining shows a higher density of collagen fibers in SKOV-3 compared to AU565 and XTM tumors in both untreated and TZM-treated tumors, suggesting that collagen fibers could act as a physical barrier that could contribute to a differential sensitivity to TZM (**Figure 3A, Mason’s Trichrome**). A higher degree of vascularity (CD31 positive cells) is observed in untreated and treated AU565 tumors compared to SKOV-3 tumors. Interestingly, AU565-derived XTM tumor shows similar vascularity to SKOV-3 tumors, suggesting that vascular bed characteristics can be altered during tumor passaging (**Figure 3B, Anti-CD31**). No significant changes were observed in the HER2 expression levels in the tumor tissue sections post-TZM treatment, although the number of HER2-positive cells is reduced in both AU565 and XTM compared to SKOV-3 tumors, indicating higher TZM toxicity in HER2 breast tumors (**Figure 3B, Anti-HER2**).

**Figure 3.**
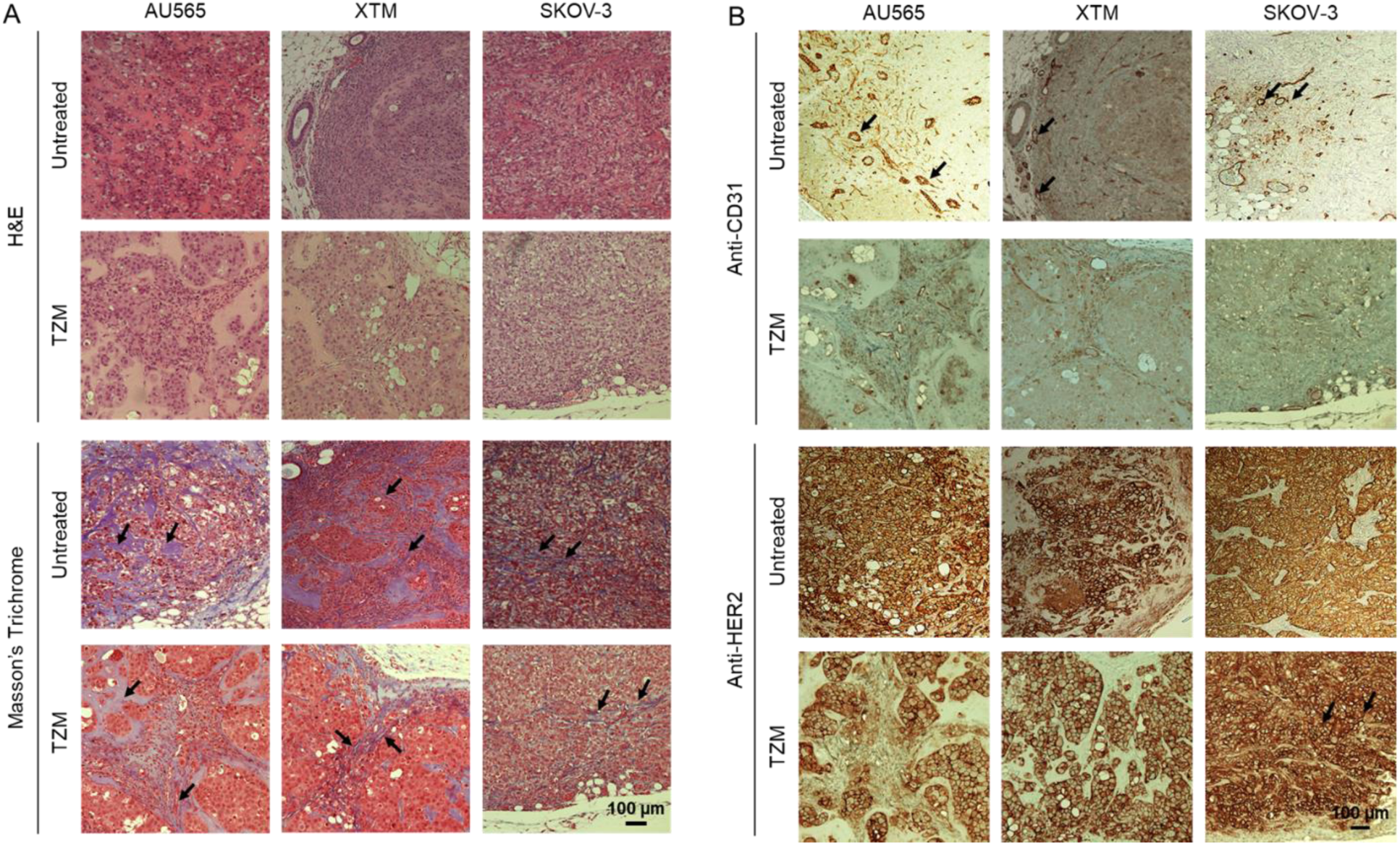
Histopathological characterization of untreated and TZM-treated (60 µg) HER2-positive AU565, XTM, and SKOV-3 tumor xenografts. Paraffinized tumor sections of tumors were processed for **(A)** H&E staining, Masson’s trichrome staining, and **(B)** anti-CD-31 and anti-HER2 immunohistochemical staining. Black arrows indicate collagen fiber bundles **(A)** and CD-31 positive cells **(B)**. NovaRED was used as peroxidase substrate (brown stain), tissue was counterstained with methyl green. Scale bar = 100 µm. Untreated AU565, n= 2; XTM, n= 3; SKOV-3, n= 2. Treated AU565, n= 5; XTM, n= 3; SKOV-3, n= 3; n= number of tumors analyzed per group.

### Quantification of TZM-HER2 engagement in human breast and ovarian tumor xenografts

Following the morphological and molecular characterization of untreated and TZM-treated HER2- positive human breast and ovarian tumor xenografts, we measured non-invasively the level of TZM-HER2 engagement in AU565 and SKOV-3 tumor xenografts (**Protocols P-1a and P-1b; Figure S1**) [16]. **Figure 4** shows analysis of (**A**) TZM-AF700/TZM-AF750 (A:D = 2:1) and (**B**) TZM-AF700/TZM-QC-1 (A:D = 3:1) mediated MFLI FRET in nude mice carrying AU565 and SKOV-3 tumor xenografts. Different A:D ratios were used to optimize the FRET assay for each NIR fluorophore FRET pair, as described previously [16]. **Figures 4A-B**, **S2-S3** indicate that in both AU565 and SKOV-3 tumors, the intensity and FRET FD% maps show an inverse relationship at 24 h and 48 h post-TZM injection. In general, the SKOV-3 tumors show relatively high intensity and low FRET signal; conversely, and the AU565 tumors display relatively low intensity and relatively higher FRET percentage (**Figure 4A**, **S2-S3**). However, the intensity maps show high variability across tumors. The quantitative pixel-by-pixel estimation of FD% levels and distribution show that AU565 tumors have a high FRET signal compared to SKOV-3 tumors using either TZM-AF700/TZM-AF750 or TZM-AF700/TZM-QC-1 FRET pairs at 24 h and 48 h post-TZM injection, indicating higher TZM-HER2 binding in AU565 compared to SKOV-3 tumors (**Figure 4C-D**). After *in vivo* MFLI imaging, the tumors were extracted and subjected to histological evaluation for TZM and HER2 tumor distribution. The IHC data shows higher TZM levels in AU565 compared to SKOV-3 tumors, thereby validating the MFLI FRET data (**Figure 5A-B).** These results suggest that SKOV-3 tumors, in comparison to AU565 tumors, have higher non-specific passive drug accumulation, whereas AU565 tumors show an elevated FRET signal due to increased TZM-HER2 drug-target engagement.

**Figure 4.**
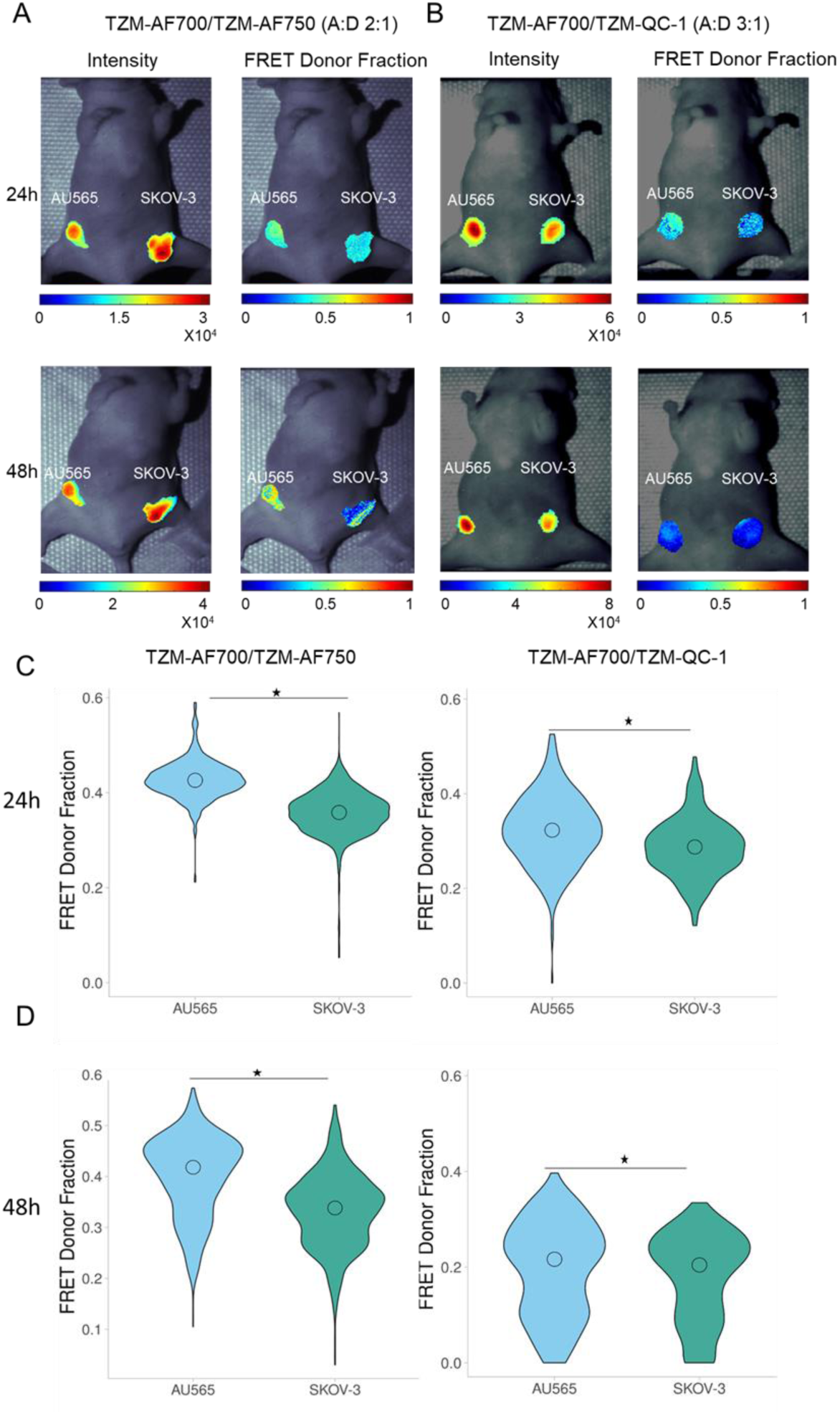
Comparative analysis of TZM-HER2 drug target engagement in labeled TZM-AF700/TZM-AF750 MFLI FRET Vs TZM-AF700/QC-1 (dark quencher) MFLI FRET using *in vivo* imaging of nude mice carrying AU565 and SKOV-3 tumor xenografts. **(A)** Mice were injected with either 20 µg TZM–AF700 and 40 µg TZM– AF750, A:D 2:1 or **(B)** 20 µg TZM–AF700 and 60 µg TZM–QC-1, A:D 3:1 and subjected to MFLI FRET imaging at 24 h and 48 h p.i. Photomicrographs show TZM donor maximum intensity (including both soluble and bound probe) and FD% maps (bound and internalized probe) for tumor ROIs in TZM–AF700/ TZM–AF750 or TZM–AF700/ TZM–QC-1 treated AU565, and SKOV-3 tumors (T), at 24 h p.i. and 48 h p.i. **(C)** Violin-plot of FD% retrieved for each tumor at 24 h p.i. and 48 h p.i. in TZM–AF700/ TZM–AF750 or (D) TZM–AF700/ TZM–QC-1 treated mice. Data presented as violin-plot indicating 25–75% pixel values, center point indicate mean with ±1.5 SD, respectively. Asterisks indicate p<0.05 (significant), *AU565 vs. SKOV-3. TZM–AF700 and TZM–AF750 treated AU565; n= 5; SKOV-3, n= 5, TZM–AF700 and TZM–QC-1 treated AU565, n= 4; SKOV-3, n= 4, n= number of tumors analyzed per group **(Supplementary Table S3-S4)**.

**Figure 5.**
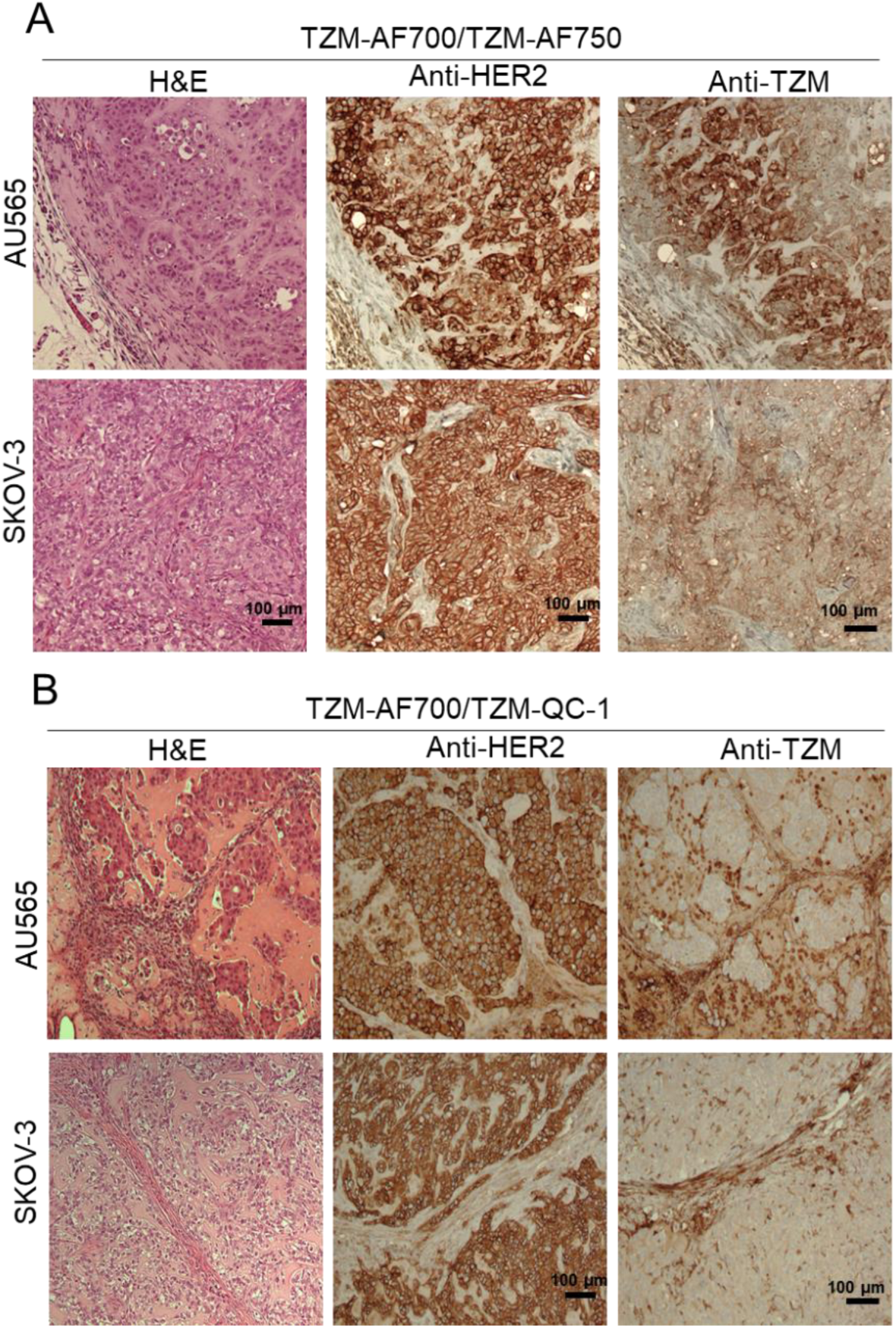
Histopathological validation of TZM-treated AU565 and SKOV-3 tumor xenografts model. Paraffinized tumor sections of tumors were processed for H&E staining, anti-HER2, and anti-TZM immunohistochemical staining in TZM–AF700/ TZM–AF750 treated mice **(A)** and TZM–AF700/ TZM–QC-1 **(B)** treated mice. NovaRED was used as peroxidase substrate (brown stain), tissue was counterstained with methyl green. Scale bar = 100 µm, n= 5 for TZM–AF700 and TZM–AF750; n=4 for TZM–AF700 and TZM– QC-1. n= number of tumors analyzed per group.

The TZM-HER2 binding was compared in XTM vs SKOV-3 tumors in nude mice injected with TZM-AF700/TZM-QC-1 (A:D 2:1) as depicted in **Protocol P-2** (**Figure S1**). The FRET data is similar in SKOV-3 across different experiments (**Figure 4 and 6**; **S2**, **S3 and S4**), strengthening the potential use of dark quencher FRET acceptors for evaluating drug-target engagement in tumor xenografts. Furthermore, we observed that HER2-positive XTM tumors display higher FD% levels, suggesting increased TZM-binding compared to HER2-positive ovarian tumors (SKOV-3) (**Figure 6B**). These results were further validated by IHC staining, indicating that breast tumors (AU565 and XTM) have higher TZM binding compared to the ovarian tumor (SKOV-3), linked to heterogeneous tumor microenvironment (**Figure 5 and 6C**). Finally, in **Figure 7A**, we used MFLI imager to measure the FRET signal in AU565 and XTM tumors to determine the TZM binding in similar tumor types (**Protocol P-3; Figure S1**). No significant difference was observed both in FD% levels and IHC validation, suggesting that tumor cells with the same origin have similar TZM binding, linked to similar TME (**Figure 7B-C**, **S5**). Taken together, we demonstrated the utility of QC-1-based dark quencher FRET for quantifying the TZM-HER2 engagement in tumors with heterogeneous TME, unraveling the potential to screen therapy-sensitive and resistant tumors at an early-stages of target characterization and identification. These results demonstrate that MFLI FRET-based measurement of drug-target engagement across different tumor systems adequately reflects the drug delivery and binding characteristics of each tumor type.

**Figure 6.**
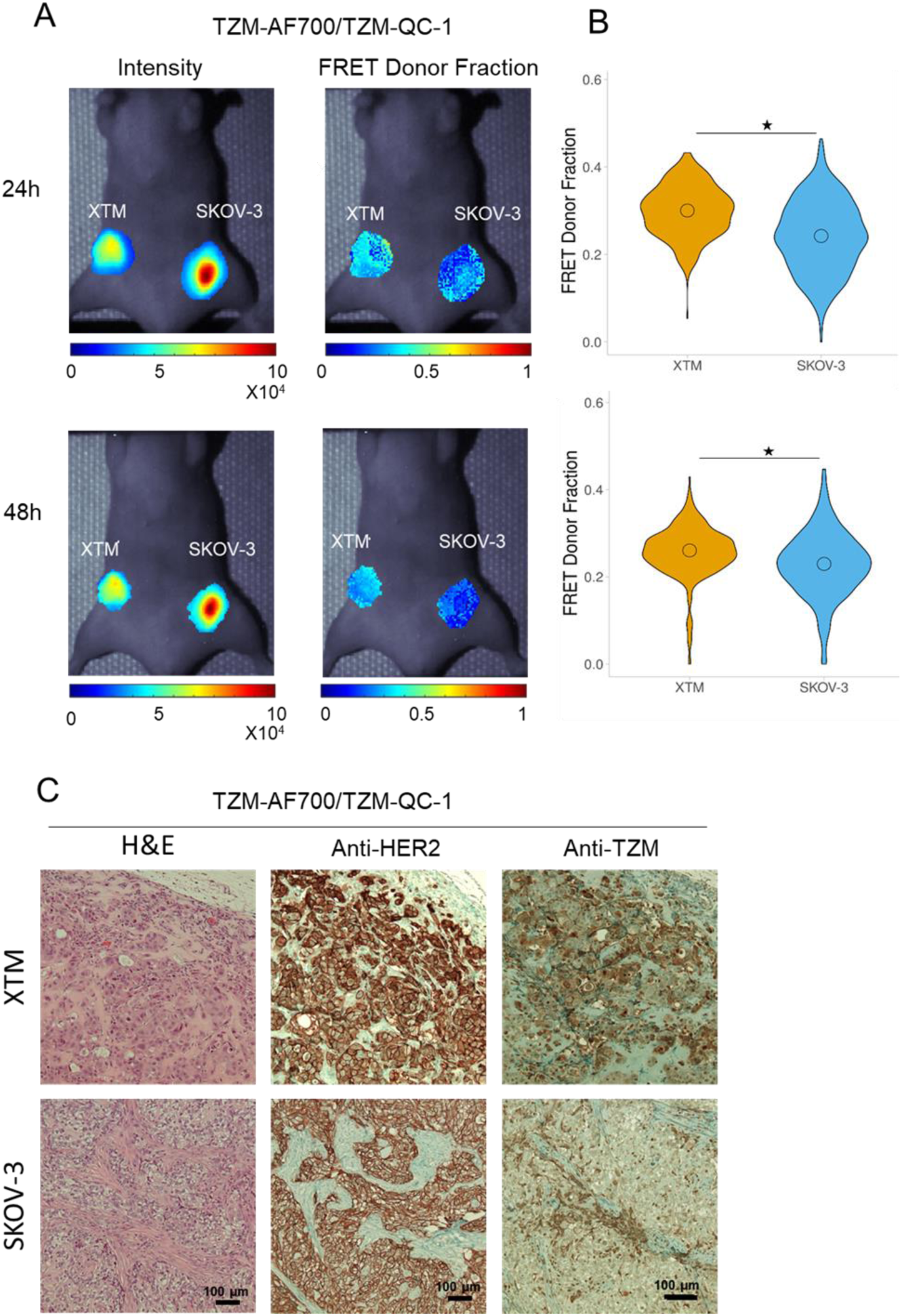
TZM-HER2 drug target engagement in labeled TZM-AF700/QC-1 (dark quencher) MFLI FRET using *in vivo* imaging in nude mice carrying XTM and SKOV-3 tumor xenografts. **(A)** Mice were injected with either 20 µg TZM–AF700 and 40 µg TZM–QC-1; A:D 2:1 and subjected to MFLI FRET imaging at 24 h and 48 h p.i. Photomicrograph show TZM donor maximum intensity (both soluble and bound probe) and FD% map (bound and internalized probe) for tumor ROIs in TZM–AF700/ TZM–QC-1 treated XTM, and SKOV-3 tumors (T), at 24 h p.i. and 48 h p.i. **(B)** Violin-plot of FD% retrieved for each tumor at 24 h p.i. and 48 h p.i. in TZM–AF700/ TZM–QC-1 treated mice. Data presented as violin-plot indicating 25–75% pixel values, center point indicate mean with ±1.5 SD, respectively. Asterisks indicate p<0.05 (significant), *XTM Vs SKOV-3; **Supplementary Table S5**. **(C)** Histopathological validation of TZM-treated XTM and SKOV-3 tumor xenografts model. Paraffinized tumor sections of tumors were processed for H&E staining, anti-HER2, and anti-TZM immuno-histochemical staining in TZM–AF700/ TZM–QC-1 treated mice. NovaRED was used as peroxidase substrate (brown stain), tissue was counterstained with methyl green. Scale bar = 100 µm, TZM–AF700 and TZM–QC-1 treated XTM, n= 3; SKOV-3, n=3. n= number of tumors analyzed per group.

**Figure 7.**
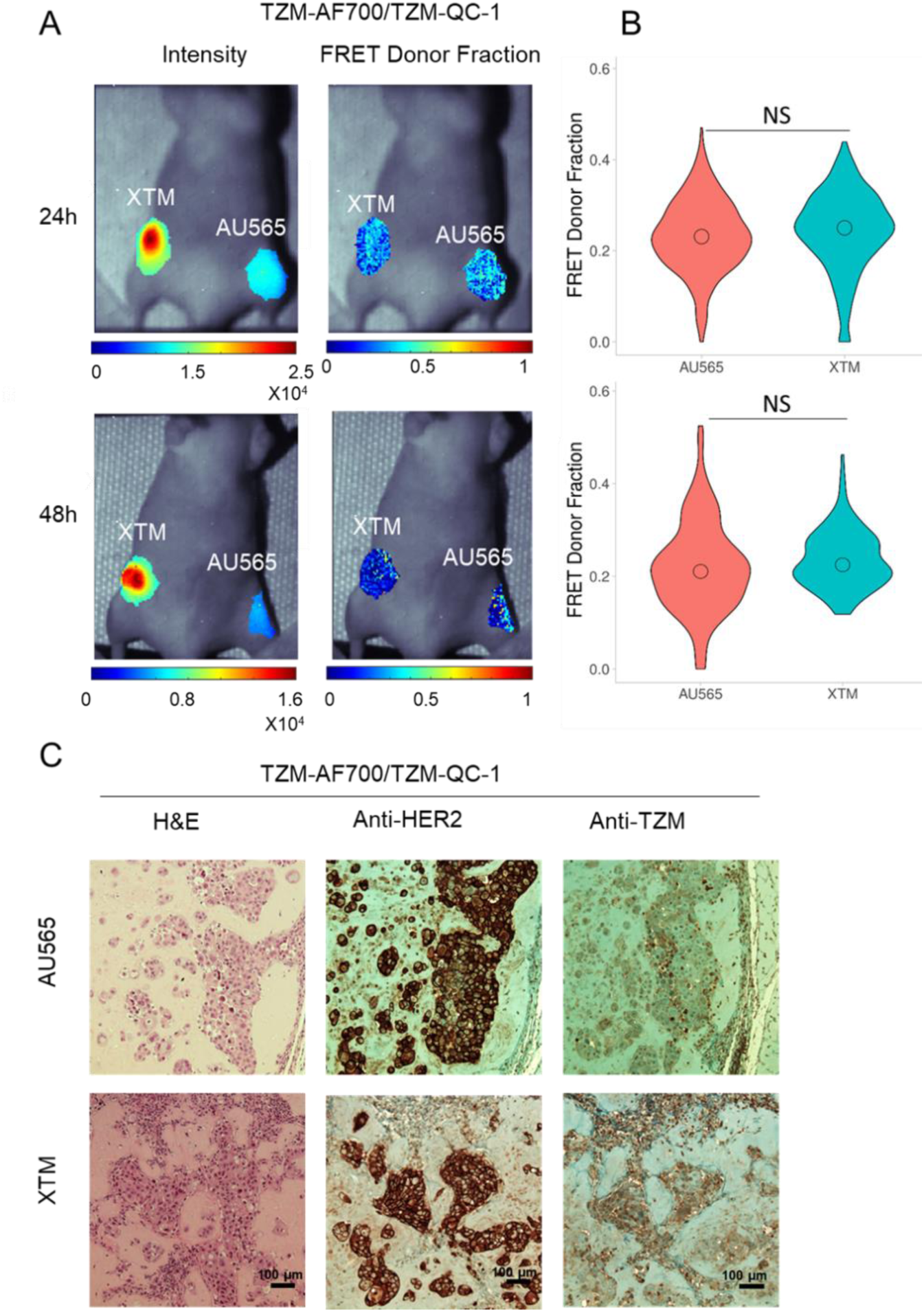
TZM-HER2 drug target engagement in labeled TZM-AF700/QC-1 (dark quencher) MFLI FRET using *in vivo* imaging in nude mice carrying AU565 and XTM tumor xenografts. **(A)** Mice were injected with either 20 µg TZM–AF700 and 60 µg TZM–QC-1; A:D 3:1 and subjected to MFLI FRET imaging at 24 h, 48 h p.i. Photomicrograph show TZM donor maximum intensity (both soluble and bound probe) and FD% map (bound and internalized probe) for tumor ROIs in TZM–AF700/ TZM–QC-1 treated AU565 and XTM tumors (T), at 24 h p.i. and 48 h p.i. **(B)** Violin-plot of FD% retrieved for each tumor at 24 h p.i. and 48 h p.i. in TZM–AF700/ TZM–QC-1 treated mice. Data presented as violin-plot indicating 25–75% pixel values, center point indicate mean with ±1.5 SD, respectively. Asterisks indicate p<0.05 (significant), *XTM Vs SKOV-3; **Supplementary Table S6**. (C) Histopathological validation of TZM-treated AU565 and XTM tumor xenografts model. Paraffinized tumor sections of tumors were processed for H&E staining, anti-HER2, and anti-TZM immuno-histochemical staining in TZM–AF700/ TZM–QC-1 treated mice. NovaRED was used as peroxidase substrate (brown stain), tissue was counterstained with methyl green. Scale bar = 100 µm, TZM–AF700 and TZM–QC-1 treated XTM, n= 3; AU565, n=3. n= number of tumors analyzed per group.

**Figure 8.**
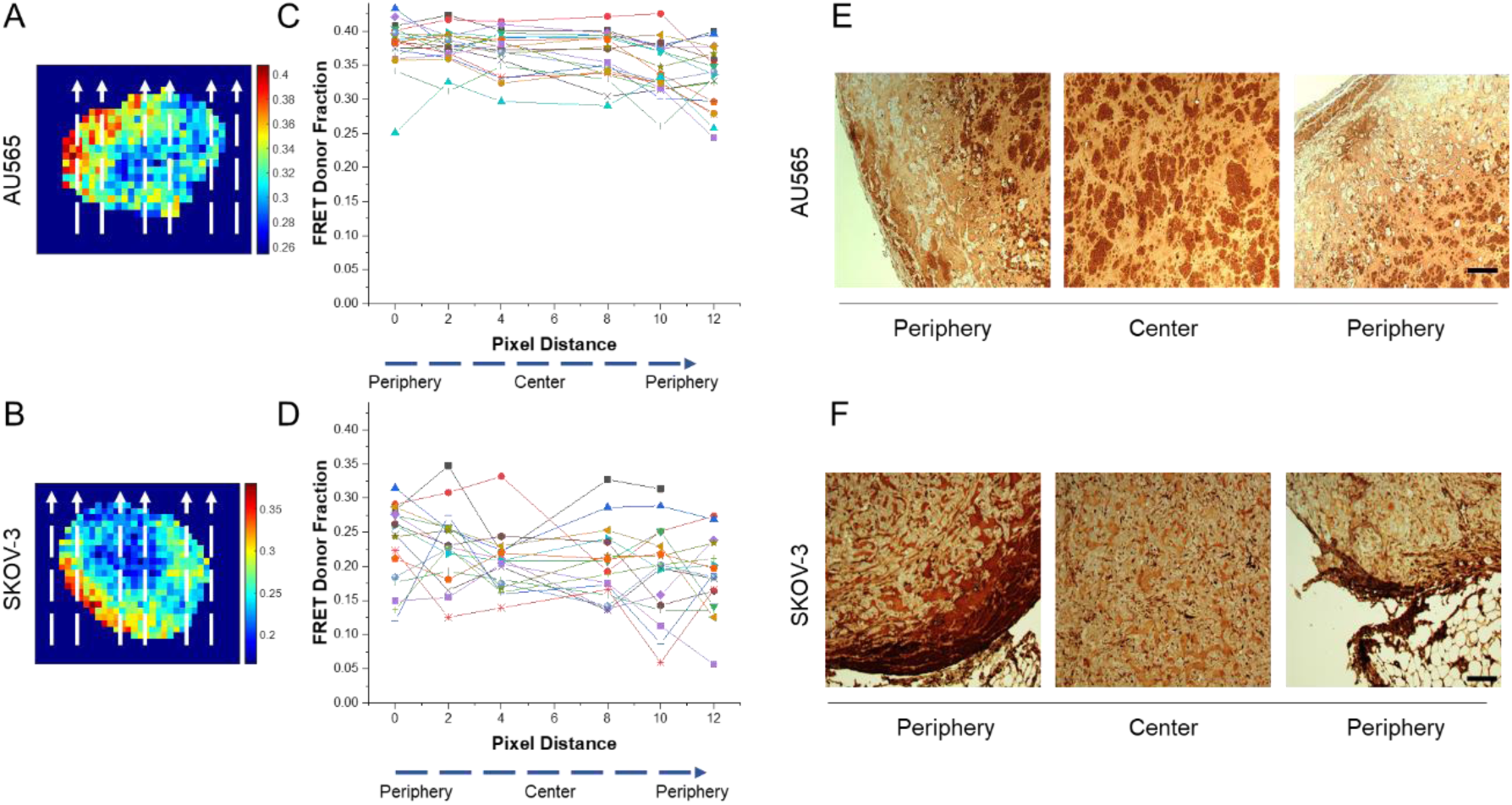
Characterization and validation of pixel-by-pixel MFLI FRET in HER2-positive AU565 and ovarian SKOV-3 tumor xenografts in nude mice carrying AU565 and SKOV-3 tumor xenografts. Pixel-by-pixel MFLI FRET map in **(A)** AU565, and **(B)** SKOV-3 tumors. Dotted lines from bottom to top indicate the pixel distribution from periphery to center to periphery. Correlation between FRET Donor Fraction (FD %) and pixel distance across tumor from periphery-to-center-to-periphery in **(C)** AU565 and **(D)** SKOV-3 tumors. Immunohistochemical validation of MFLI FRET in **(E)** AU565, and **(F)** SKOV-3 tumors was performed using anti-TZM primary antibody. NovaRED was used as peroxidase substrate (brown color), sections were counterstained with methyl green. Scale bar = 40 μm.

### Intratumoral distribution of TZM in HER2-positive human breast and ovarian tumor xenografts in mice

The inability of TZM to distribute across different tumor regions even in the presence of elevated HER2 expression is one of the leading causes of failure of TZM therapy. We posit that the inability of TZM to bind HER2-expressing tumor cells can be associated with variations in TME as shown here in different types of HER2-positive tumor xenografts (AU565 vs SKOV-3). We analyzed the pixel-by-pixel MFLI FRET mapping in AU565 and SKOV-3 tumors within the tumor center and periphery and correlated the FD% fraction with pixel distance, indicated in dashed arrows (**Figure 8A-D**). The data suggest that the FD% fraction is distributed uniformly in AU565 cells across the pixel distance from the periphery to the center of tumors. In contrast, in SKOV-3 the FD% fraction is higher towards the periphery and reduced closer to the tumor center. A similar pattern is observed in IHC analysis, where unvarying anti-TZM staining is observed across the AU565 tumor, while in SKOV-3 anti-TZM staining is restricted to the periphery, thereby validating the intratumoral distribution of TZM observed in MFLI FRET imaging map (**Figure 8E-F**). These results suggest that MFLI FRET imaging can be used to non-invasively quantify the drug-target engagement in tumor xenografts as well as characterize the intratumoral spatial distribution of TZM-HER2 drug-target engagements across live and intact tumor xenografts. In addition, the distribution of TZM appears to be primarily limited to the peripheral region of the less TZM-sensitive tumors (SKOV-3). TZM’s inability to access the HER2-expressing cells in the central tumor region may be due to collagen-mediated stiffness and reduced vascularity characteristic of SKOV-3 tumor xenografts.

## CONCLUSION

We present dark quencher-based MFLI FRET as a unique noninvasive approach to measure the drug-target engagement of HER2-positive human breast and ovarian tumor xenografts mice model, which vary in their TME. Our study demonstrates dark quencher based MFLI FRET as a direct measure of tumor heterogeneity/ resistance linked to high collagen content, vascularity, and drug distribution. This molecular imaging approach is expected to enable the development of new targeted drugs and/or interventions for improved drug delivery efficacy, facilitating the therapeutic management of breast cancer.

## ACKNOWLEDGMENTS

We would like to acknowledge the support of the AMC imaging core for the use of the LSM 880 confocal microscope and help with the collection of NIR FLIM FRET microscopy data. This work was funded by the National Institutes of Health grants R01 CA250636, R01 CA207725, and R01 CA237267 and R01 CA271371.

## AUTHOR’S CONTRIBUTION

M.B. & X.I conceived the original idea. M.B. & X.I. designed the research study. C.S, & M.B acquired and analyzed the NIR FLIM microscopy data sets. AR and JTS acquired the MFLI data sets. A.V, V.P, & J.T.S. processed the MFLI data sets. A.V, V.P, & J.T.S. performed the data processing and analyses of results. A.V, V.P, M.B & X.I interpreted the results. All authors contributed to the writing of the manuscript. All authors have approved the final version of the manuscript.

## SUPPLEMENTARY FIGURES

**Figure S1.**
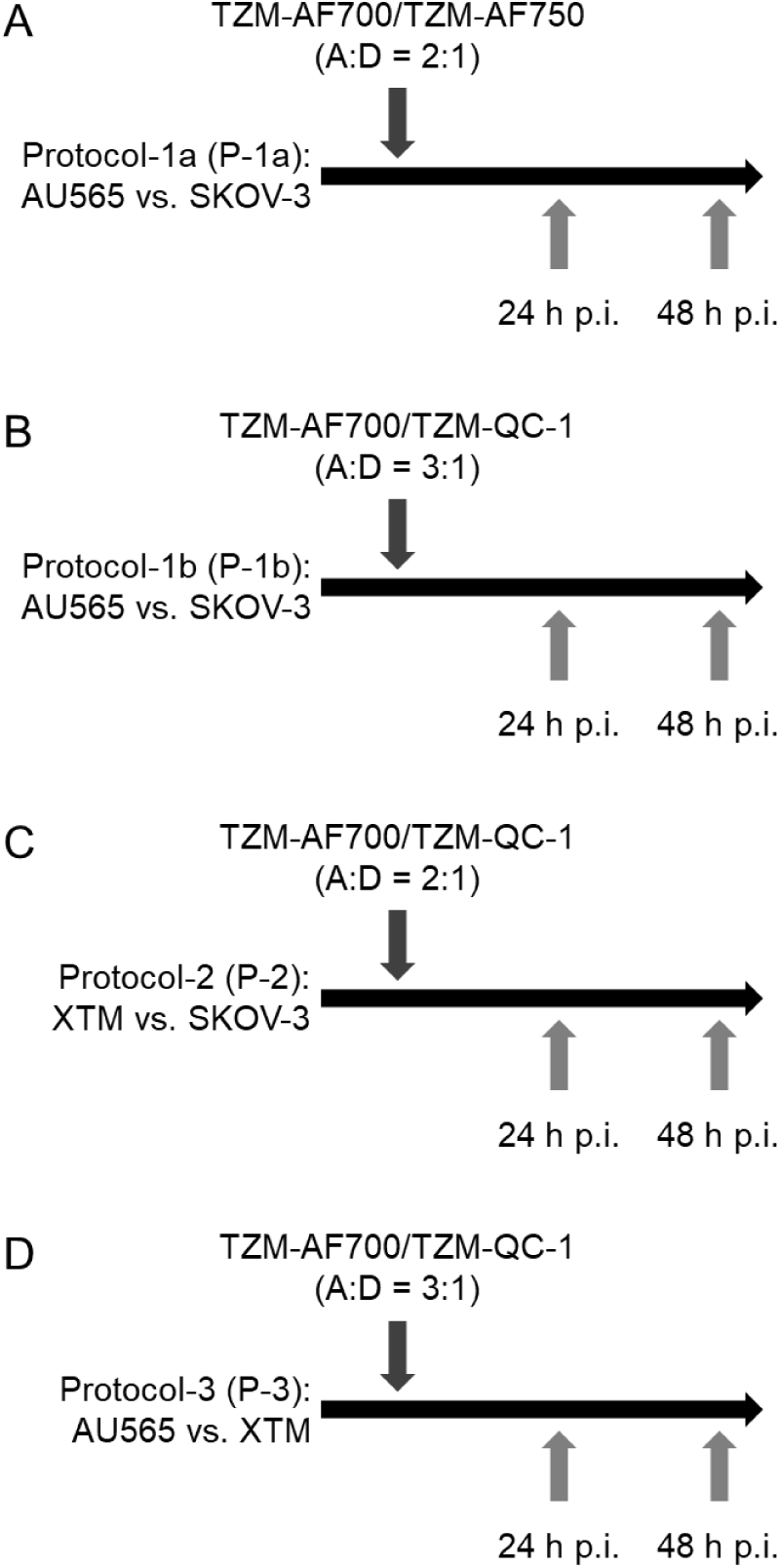
Injection and *in vivo* imaging protocols for AU565, XTM, and SKOV-3 tumor xenografts-bearing mice: **(A) Protocol 1a (P-1a)**, mice bearing AU565 and SKOV-3 tumor xenografts were injected intravenously with TZM-AF700/TZM-AF750 (A:D 2:1) and live intact animal MFLI data was captured at 24 h and 48 h p.i.; **(B) Protocol-1b (P-1b),** mice bearing AU565 and SKOV-3 tumor xenografts were injected intravenously with TZM-AF700/TZM-QC-1 (A:D 3:1) and live intact animal MFLI data was captured at 24 h and 48 h p.i.; **(C) Protocol 2 (P-2)**, mice bearing XTM and SKOV-3 tumor xenografts were injected intravenously with TZM-AF700/TZM-QC-1 (A:D 2:1) and live intact animal MFLI data was captured at 24 h and 48 h p.i.; **(D) Protocol-3 (P-3)**, mice bearing AU565 and XTM tumor xenografts were injected intravenously with TZM-AF700/TZM-QC-1 (A:D 3:1) and live intact animal MFLI data was captured at 24 h and 48 h p.i..

**Figure S2.**
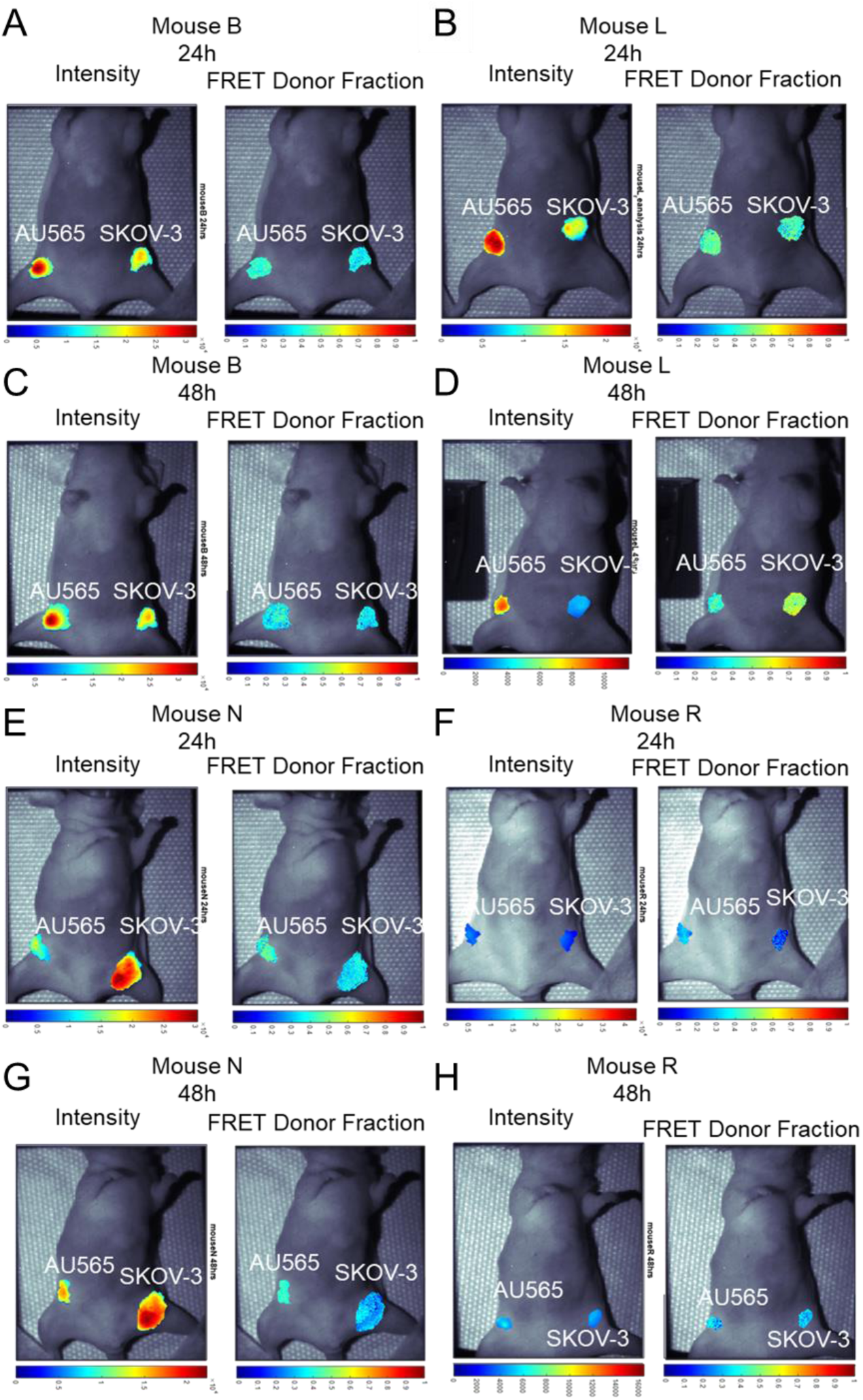
MFLI FRET *in vivo* imaging in nude mice carrying AU565 and SKOV-3 tumor xenografts (Protocol P-1a). **(A-H)** Photomicrographs show TZM donor maximum intensity ROIs (both soluble and bound probe) and FD% map (bound and internalized probe) in TZM–AF700/ TZM–AF750 treated AU565 and SKOV-3 tumors (T), at 24 h p.i. (**A-B, E-F**) and 48 h p.i. (**C-D and G-H**); TZM–AF700 and TZM–AF750 treated AU565, n= 5; SKOV-3, n= 5, n= number of tumors analyzed per group.

**Figure S3.**
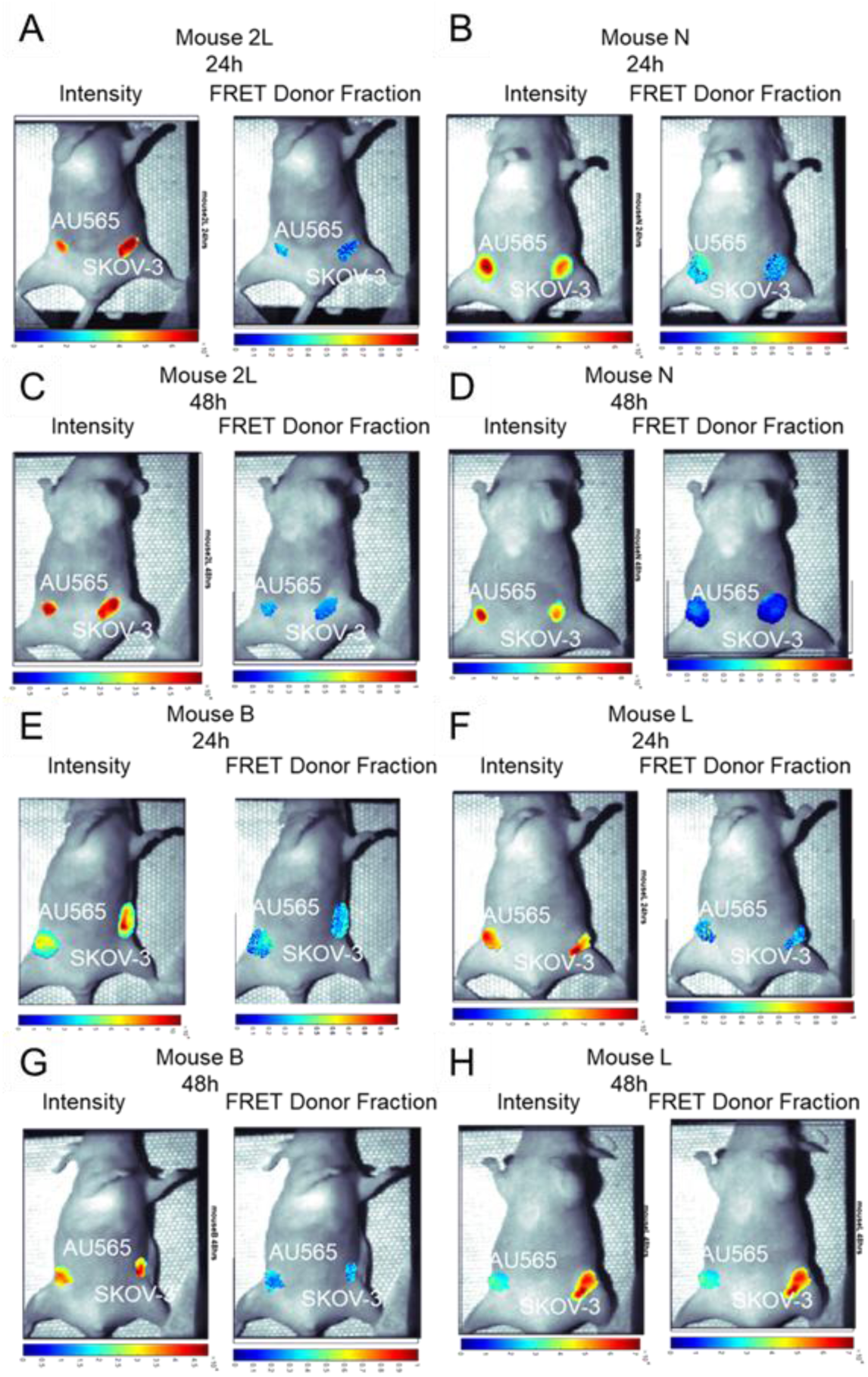
MFLI FRET *in vivo* imaging in nude mice carrying AU565 and SKOV-3 tumor xenografts (Protocol P-1b). **(A-H)** Photomicrographs show TZM donor maximum intensity ROIs (both soluble and bound probe) and FD% map (bound and internalized probe) in TZM–AF700/ TZM–QC-1 treated AU565 and SKOV-3 tumors (T), at 24 h p.i. (**A-B and E-F**) and 48 h p.i. (**C-D and G-H**); TZM–AF700 and TZM–QC-1 treated AU565, n= 4; SKOV-3, n= 4, n= number of tumors analyzed per group.

**Figure S4.**
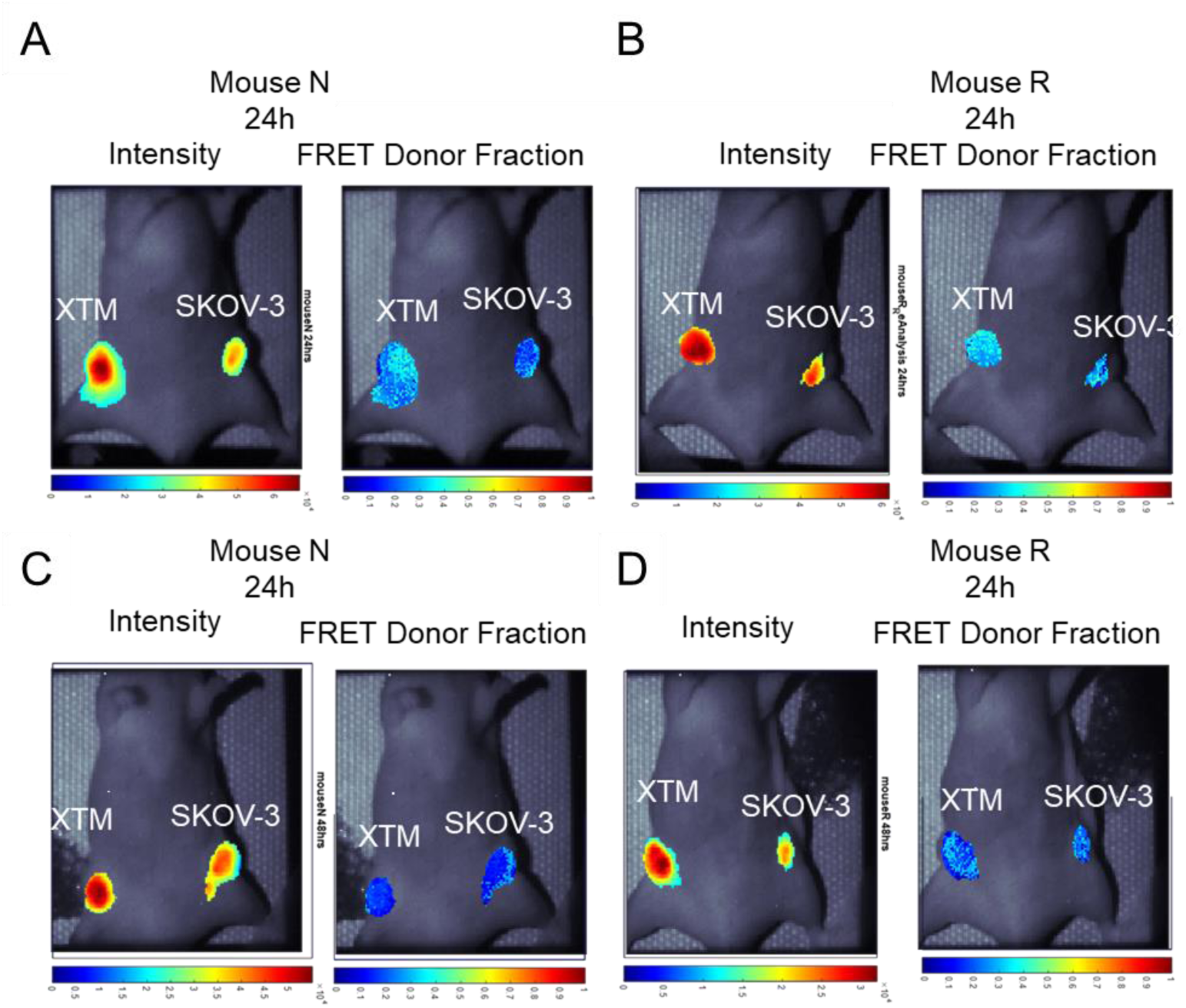
MFLI FRET *in vivo* imaging in nude mice carrying XTM and SKOV-3 tumor xenografts (Protocol P-2). **(A-D)** Photomicrographs show TZM donor maximum intensity (both soluble and bound probe) and FD% map (bound and internalized probe) for tumor ROIs in TZM– AF700/ TZM–QC-1 treated XTM and SKOV-3 tumors (T), at 24 h p.i. **(A-B)** and 48 h **(C-D)** p.i. TZM–AF700 and TZM–QC-1 treated XTM, n= 3; SKOV-3, n=3. n= number of tumors analyzed per group.

**Figure S5.**
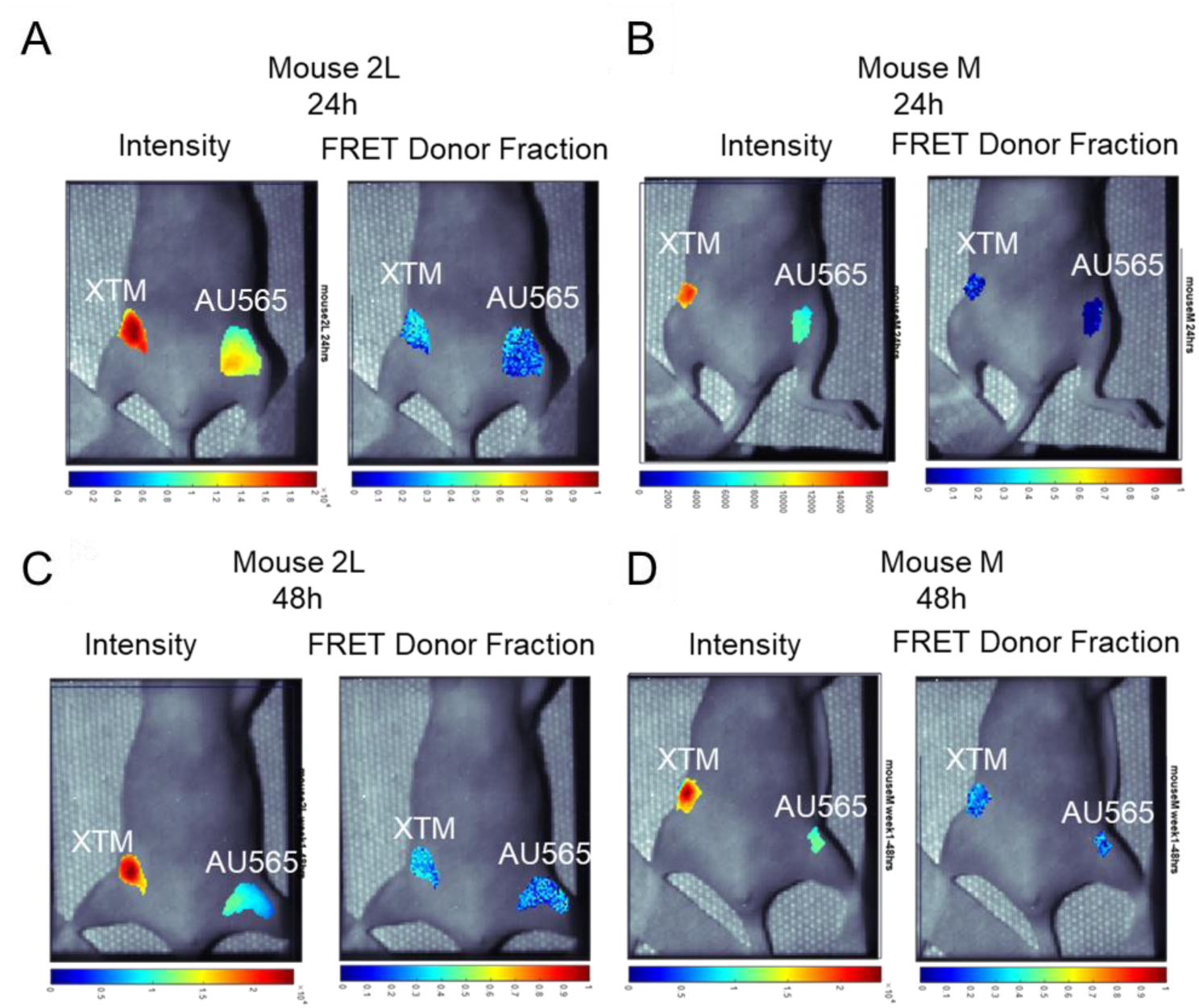
MFLI FRET *in vivo* imaging in nude mice carrying AU565 and XTM tumor xenografts (Protocol P-3). **(A-B)** Photomicrographs show TZM donor maximum intensity (both soluble and bound probe) and FD% map (bound and internalized probe) for tumor ROIs in TZM–AF700/ TZM–QC-1 treated AU565 and XTM tumors (T), at 24 h p.i. (**A-B**) and 48 h p.i. (**C-D**); TZM– AF700 and TZM–QC-1 treated XTM, n= 3; AU565, n=3. n= number of tumors analyzed per group.

**Supplementary table S1.** Experimental reagents used.

| Product | Company | Catalogue No |
| --- | --- | --- |
| Amicon Ultra-4-centrifugal filter units (MWCO 30 kDa) | Sigma Aldrich, St. Louis, MO | Z648035 |
| Anti-HER2 primary antibody | Thermo Fisher Scientific Inc, Waltham, MA | MA5-12759 |
| Trastuzumab (Anti-Human HER2, Humanized Antibody) | MedChemExpress, NJ | HY-P9907 |
| HER2/ErbB2 (29D8) Rabbit mAb | Cell Signaling, Technology | 2165 |
| Monoclonal rabbit CD31 | Cell Signaling Technology, Inc., Danvers, MA | 77699 |
| Rabbit monoclonal anti-TZM | R&D Systems, Bio-Techne, Minneapolis, MN | MAB95471-100 |
| Goat anti-mouse labeled with Alexa Fluor 488 | Abcam Inc | ab181289 |
| Alexa Fluor 700 NHS ester | Thermo Fisher Scientific Inc, Waltham, MA | A20110 |
| Alexa Fluor 750 NHS ester | Thermo Fisher Scientific Inc, Waltham, MA | A37575 |
| IRDye® QC-1 NHS Ester | Lincoln, NB, USA | 929-70030 |
| Olympus BX40 microscope equipped with an Infinity 3 camera | Lumenera Inc., Ottawa, ON, Canada | BX40-B |
| BME001-05 | R&D Systems Inc, Minneapolis, MN, USA | BME001-01 |
| Nude mice, CrTac: NCr-Foxn1nu | Taconic Biosciences, Rensselaer, NY | NCRNU-F |
| Vectastain ABC Elite kit | Vector Labs, Burlingame, CA | PK-6100 |
| Vectro NovaRED | Vector Labs, Burlingame, CA | SK-4800 |
| Methyl Green | Sigma Aldrich, St. Louis, MO | M8884 |
| AU565 | ATCC, Manassa, VA | CRL-2351 |
| SKOV-3 | ATCC, Manassa, VA | HTB-77 |
| RPMI 1640 | Thermo Fisher Scientific Inc, Waltham, MA | 11875093 |
| McCoy's media | Thermo Fisher Scientific Inc, Waltham, MA | 16600082 |
| Hanks' Balanced Salt Solution (HBSS) | Thermo Fisher Scientific Inc, Waltham, MA | 88284 |
| μ-Slide 8 Well | Ibidi, Fitchburg, WI, | 80826 |
| Fetal bovine serum | ATCC, Manassa, VA | 30-2021 |
| HEPES | Thermo Fisher Scientific Inc, Waltham, MA, | 15630080 |
| Penicillin/Streptomycin | Thermo Fisher Scientific Inc, Waltham, MA, | 15070063 |

**Supplementary table S1. Contd.**
|  |  |  |
| --- | --- | --- |
| Beckman DU-640 Spectrophotometer | GMI | SKU: 8043-30-1090 |
| Digital micromirror (DMD) device | Texas Instruments, TX | DLi 4110 |
| Titanium/ Sapphire laser, Chameleon Ultra II | Coherent, Inc., Santa Clara, CA |  |
| Mai Tai Ultrafast Laser | Spectra-Physics, CA |  |
| SPCImage NG Data Analysis Software | Becker & Hickl GmbH, Berlin, Germany |  |
| Intensified CCD (ICCD) camera | Picostar HR, Lasision GmbH, Bielefeld, Germany |  |

**Supplementary Table S2.**
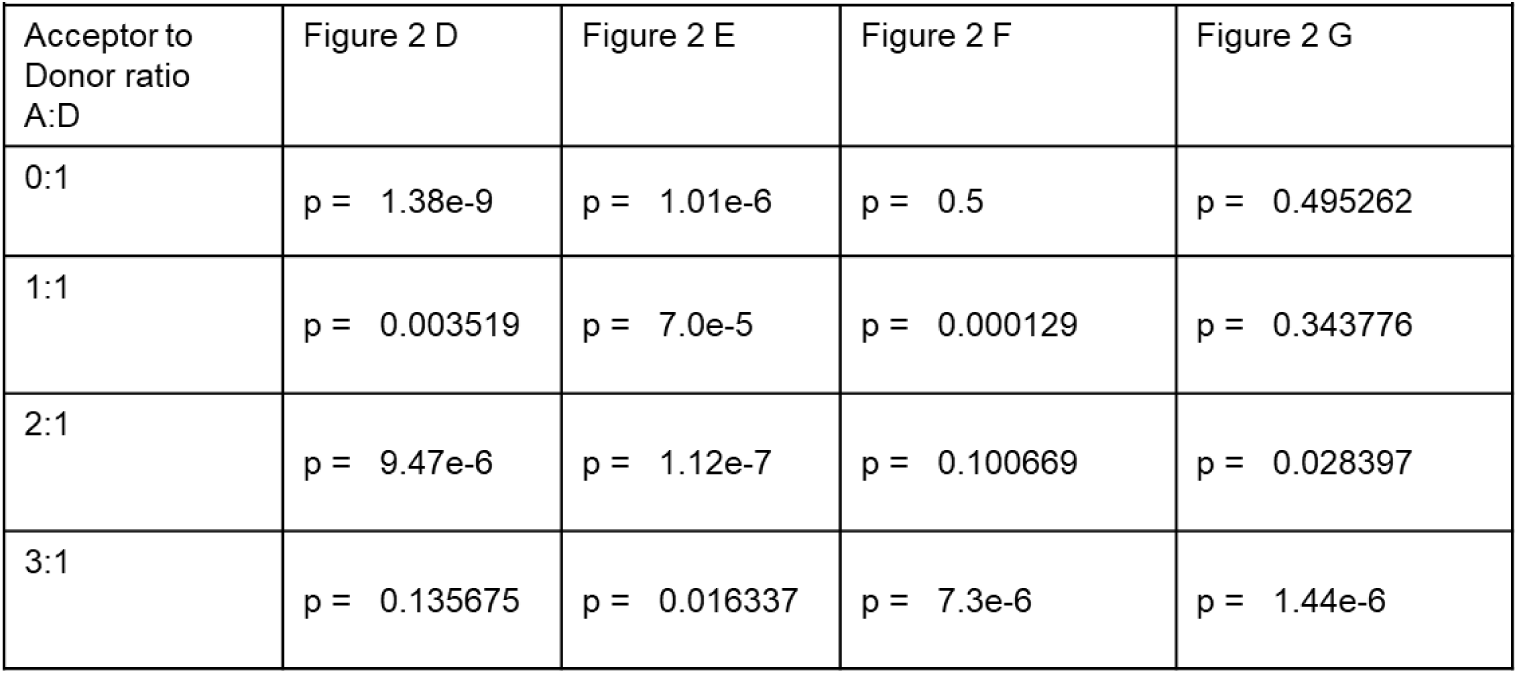
Statistical analysis for data presented in Figure 2 D-G.

**Supplementary Table 3.** Statistical analysis for data presented in Figure 4 C-D.

Supplementary Table S3 and S4. Statistical analysis for data presented in Figure 4 C-D
|  |  |  |  |  |  |  |  |
| --- | --- | --- | --- | --- | --- | --- | --- |
| TzM-AF700/TzM-AF750<br>24 h |  |  |  | TzM-AF700/TzM-AF750<br>48 h |  |  |  |
|  | N total<br>(pixels) | Mean | Standard<br>Deviation |  | N total<br>(pixels) | Mean | Standard<br>Deviation |
| AU565 | 360 | 0.4276 | 0.040045 | AU565 | 1316 | 0.4015 | 0.0779 |
| SKOV-3 | 1638 | 0.357375 | 0.047042 | SKOV-3 | 1826 | 0.3331 | 0.0727 |
| Two sample T-TEST; p-value 5.7E-261; p-value $\leq 0.05$ is significant. | | | | Two sample T-TEST; p-value 6E-261; p-value $\leq 0.05$ is significant. | | | |

**Supplementary Table 4.**
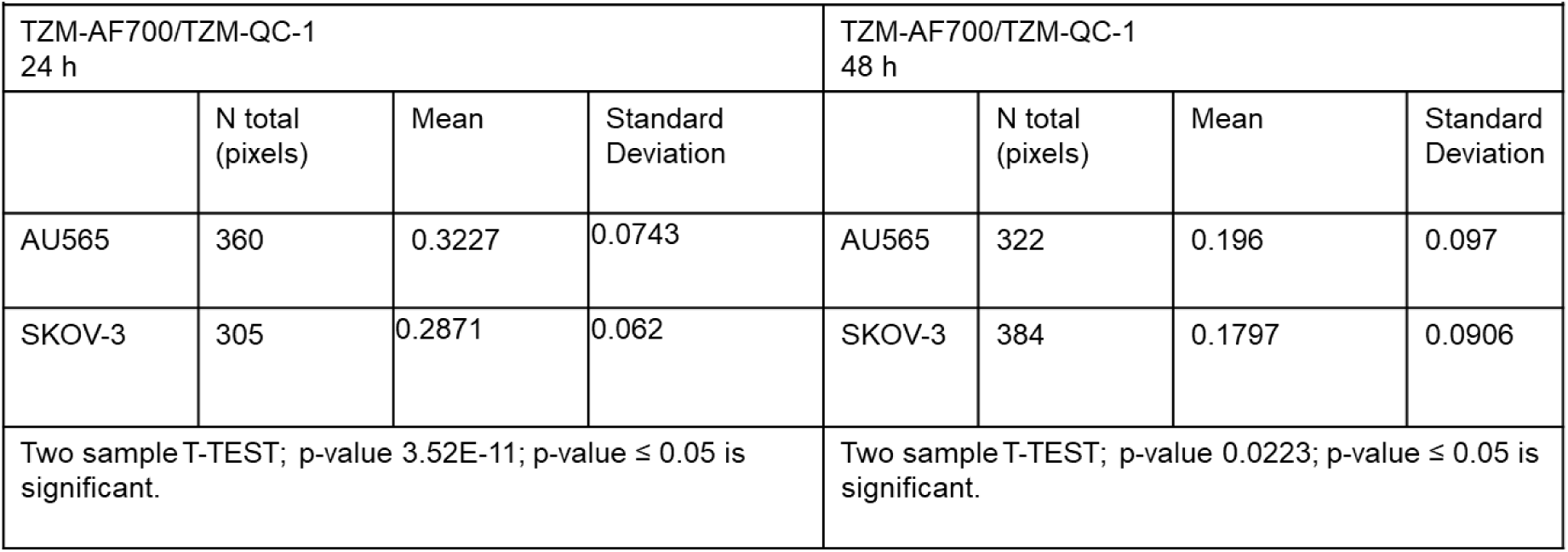
Statistical analysis for data presented in Figure 4 C-D.

|  |  |  |  |  |  |  |  |
| --- | --- | --- | --- | --- | --- | --- | --- |
| TzM-AF700/TzM-QC-1<br>24 h |  |  |  | TzM-AF700/TzM-QC-1<br>48 h |  |  |  |
|  | N total<br>(pixels) | Mean | Standard<br>Deviation |  | N total<br>(pixels) | Mean | Standard<br>Deviation |
| AU565 | 360 | 0.3227 | 0.0743 | AU565 | 322 | 0.196 | 0.097 |
| SKOV-3 | 305 | 0.2871 | 0.062 | SKOV-3 | 384 | 0.1797 | 0.0906 |
| Two sample T-TEST; p-value 3.52E-11; p-value $\leq 0.05$ is significant. | | | | Two sample T-TEST; p-value 0.0223; p-value $\leq 0.05$ is significant. | | | |

**Supplementary Table S5.**
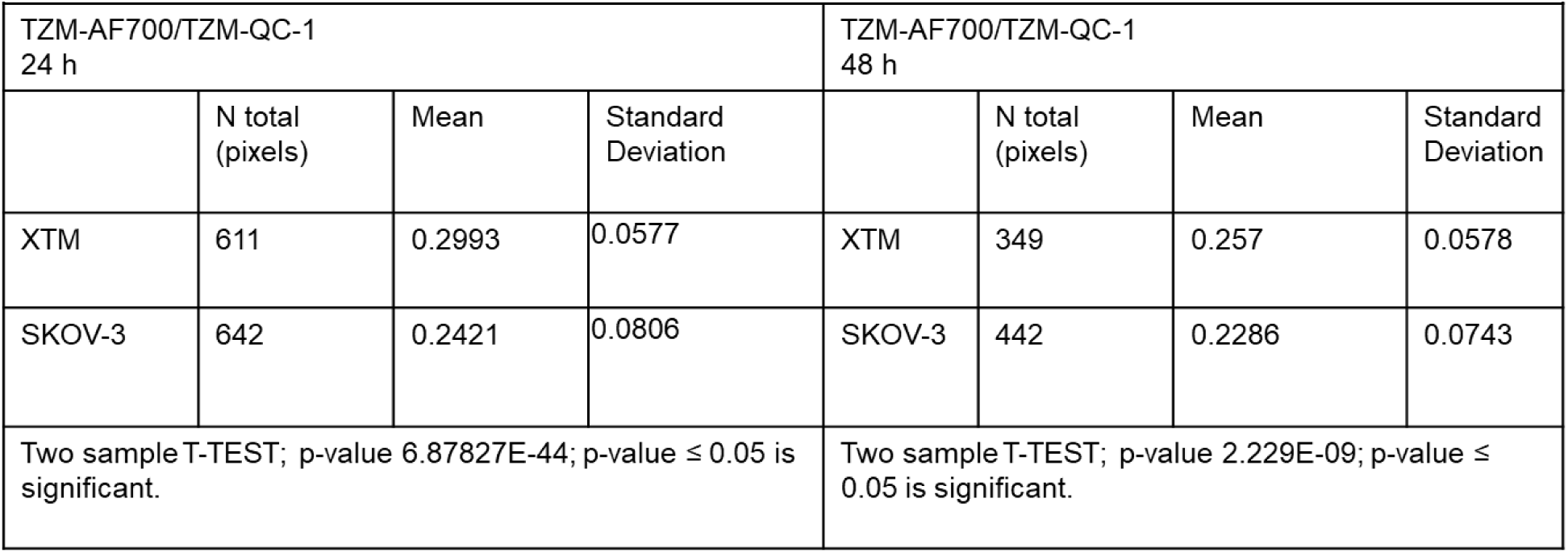
Statistical analysis for data presented in Figure 6 B.

| Supplementary Table 5. Statistical analysis for data presented in Figure 6 B |  |  |  |  |  |  |  |
| --- | --- | --- | --- | --- | --- | --- | --- |
| TQM-AF700/TQM-QC-1<br>24 h |  |  |  | TQM-AF700/TQM-QC-1<br>48 h |  |  |  |
|  | N total<br>(pixels) | Mean | Standard<br>Deviation |  | N total<br>(pixels) | Mean | Standard<br>Deviation |
| XTM | 611 | 0.2993 | 0.0577 | XTM | 349 | 0.257 | 0.0578 |
| SKOV-3 | 642 | 0.2421 | 0.0806 | SKOV-3 | 442 | 0.2286 | 0.0743 |
| Two sample T-TEST; p-value 6.87827E-44; p-value $\leq 0.05$ is significant. | | | | Two sample T-TEST; p-value 2.229E-09; p-value $\leq 0.05$ is significant. | | | |

**Supplementary Table S6.** Statistical analysis for data presented in Figure 7 B.

| Supplementary Table 6. Statistical analysis for data presented in Figure 7 B |  |  |  |  |  |  |  |
| --- | --- | --- | --- | --- | --- | --- | --- |
| TQM-AF700/TQM-QC-1<br>24 h |  |  |  | TQM-AF700/TQM-QC-1<br>48 h |  |  |  |
|  | N total<br>(pixels) | Mean | Standard<br>Deviation |  | N total<br>(pixels) | Mean | Standard<br>Deviation |
| AU565 | 483 | 2.33E-01 | 0.07919 | AU565 | 151 | 0.2202 | 0.0966 |
| XTM | 487 | 2.43E-01 | 0.0876 | XTM | 151 | 0.2362 | 0.0626 |
| Two sample T-TEST; p-value 0.0707; p-value $\leq 0.05$ is significant. | | | | Two sample T-TEST; p-value 0.0871; p-value $\leq 0.05$ is significant. | | | |

